# Membrane insertion of α-xenorhabdolysin in near-atomic detail

**DOI:** 10.1101/312314

**Authors:** Evelyn Schubert, Ingrid R. Vetter, Daniel Prumbaum, Pawel A. Penczek, Stefan Raunser

## Abstract

α-Xenorhabdolysins (Xax) are α-pore-forming toxins (α-PFT) from pathogenic bacteria that form 1-1.3 MDa large pore complexes to perforate the host cell membrane. PFTs are used by a variety of bacterial pathogens as an offensive or defensive mechanism to attack host cells. Due to the lack of structural information, the molecular mechanism of action of Xax toxins is poorly understood. Here, we report the cryo-EM structure of the XaxAB pore complex from *Xenorhabdus nematophila* at an average resolution of 4.0 Å and the crystal structures of the soluble monomers of XaxA and XaxB at 2.5 Å and 3.4 Å, respectively. The structures reveal that XaxA and XaxB are built similarly and appear as heterodimers in the 12-15 subunits containing pore. The structure of the XaxAB pore represents therefore the first structure of a bi-component α-PFT. Major conformational changes in XaxB, including the swinging out of an amphipathic helix are responsible for membrane insertion. XaxA acts as an activator and stabilizer for XaxB that forms the actual transmembrane pore. Based on our results, we propose a novel structural model for the mechanism of action of Xax toxins.

## INTRODUCTION

Pore-forming toxins (PFTs) are soluble proteins produced by bacteria and higher eukaryotes, that spontaneously form pores in biomembranes and act as toxins [1]. Dependent on their transmembrane region, which is formed either by α-helices or β-strands, PFTs are classified as α-PFTs and β-PFTs [2, 3]. A common trait of all PFTs is the conversion from a soluble monomer to a membrane-embedded oligomer [3], however, a different mechanism has been recently found for ABC toxins [4]. Specific targeting of the PFTs to the host membrane involves mostly recognition of specific proteins, glycans or lipids on the target membrane. Conformational changes resulting in the oligomerization and membrane perforation are triggered by receptor binding, catalytic cleavage, pH change or other factors [2]. The sequential order of oligomerization and membrane penetration including the formation of an oligomeric prepore is still a matter of debate [5]. The size of the oligomers ranges from tetrameric pores in Cry1Aa [6] and heptameric pores in the anthrax protective antigen [7] to 30-50-meric pores in cholesterol-dependent cytolysins (CDCs) [1, 8].

PFTs can be further divided into two groups [9]. PFTs of the first group perforate membranes by forming stable pores resulting in an uncontrolled influx and efflux of ions and other biomolecules. This destroys ion gradients and electrochemical gradients at the membrane. The toxins of the second group also perforate the membrane, but use the transmembrane channel to specifically translocate toxic enzymes into the host. Binary toxins, also called AB toxins [10] and also recently characterized ABC toxins [11] belong to the latter group. A prominent AB toxin is the anthrax toxin [12], where component B, the protective antigen, forms a translocation pore through which lethal factor or edema factor, different A components, are translocated.

α-Xenorhabdolysin is a PFT that has been first isolated from the bacterium *Xenorhabdus nematophila* [13]. Xenorhabdolysins are also found in other entomopathogenic bacteria, such as *Photorhabdus luminescens*, and human pathogenic bacteria, such as *Yersinia enterocolitica* and *Proteus mirabilis* [14]. They are composed of two subunits, namely XaxA (45 kDa) and XaxB (40 kDa) and are only active when the two components act together [14]. Xenorhabdolysins, that were suggested to be binary toxins [14, 15], perforate the membranes of erythrocytes, insect granulocytes and phagocytes and induce apoptosis [14, 15]. The mechanism of action of xenorhabdolysins including the interaction between component A and B, oligomerization and pore formation has remained enigmatic so far.

Structural prediction using the PHYRE2 server [16] does not yield any significant similarities for XaxB. XaxA cytotoxins, however, are predicted to be similar to two pore-forming cytolysins, Cry6Aa from *Bacillus thuringiensis* [17, 18] and binding component B of hemolysin BL (Hbl-B) from *Bacillus cereus* [19]. The best characterized cytolysin is probably ClyA from *Escherichia coli* and *Salmonella enterica* strains. The structure of ClyA has been determined in its soluble form [20] and pore conformation [21] and the mechanism of pore formation mechanism has been extensively studied [22]. However, in contrast to XaxAB, ClyA only contains one component. Thus, despite the structural similarity, the mechanism of action must be different.

So far structural data on xenorhabdolysins are missing limiting our understanding of these important type of toxins. Here, we used a hybrid structural biology approach combining X-ray crystallography and electron cryomicroscopy (cryo-EM) to determine the crystal structures of XaxA and XaxB from *Xenorhabdus nematophila* as soluble monomers and the cryo-EM structure of the XaxAB pore complex.

## RESULTS & DISCUSSION

### Structure of XaxA and XaxB soluble monomers

In two different experiments, we independently expressed and purified XaxA and XaxB (Methods). The protein quantity and quality of both proteins was sufficient (Figure S1a-d) to perform crystallization experiments. We obtained well diffracting crystals of both XaxA and XaxB in their soluble monomeric form and solved their structures to 2.5 Å and 3.4 Å, respectively (Figure 1a-b, Table S1).

Both XaxA and XaxB have a long rod-shaped structure and are mainly composed of α-helices (Figure 1a-b). XaxA and XaxB have a similar domain organization. They both contain a tail domain that is formed by a five-helix bundle (αA, αB, αC, αG and αH) and elongated neck and head domains. The five-helix bundle motif has so far only been described for ClyA and ClyA-type toxins [22]. Like in the case of ClyA-type toxins the N-terminal helices (αA) of XaxA and XaxB are significantly shorter than in ClyA, where it plays a crucial role in pore formation (Figure S2a). Interestingly, XaxA contains two large loops connecting the helices, a big hook-shaped loop (aa 136-169) between helices αB and αC at the top of the tail domain and an additional loop (aa 202-215) dividing helix αC (Figure 1a). The four XaxB molecules in the asymmetric unit differed considerably in their tail domain (Figure S3). Especially, helices αB and αC that protrude slightly from the five-helix-bundle take different positions. Although this might be due to tight crystal packing, it also indicates a certain degree of flexibility of the tail domain of XaxB.

**Figure 1.**
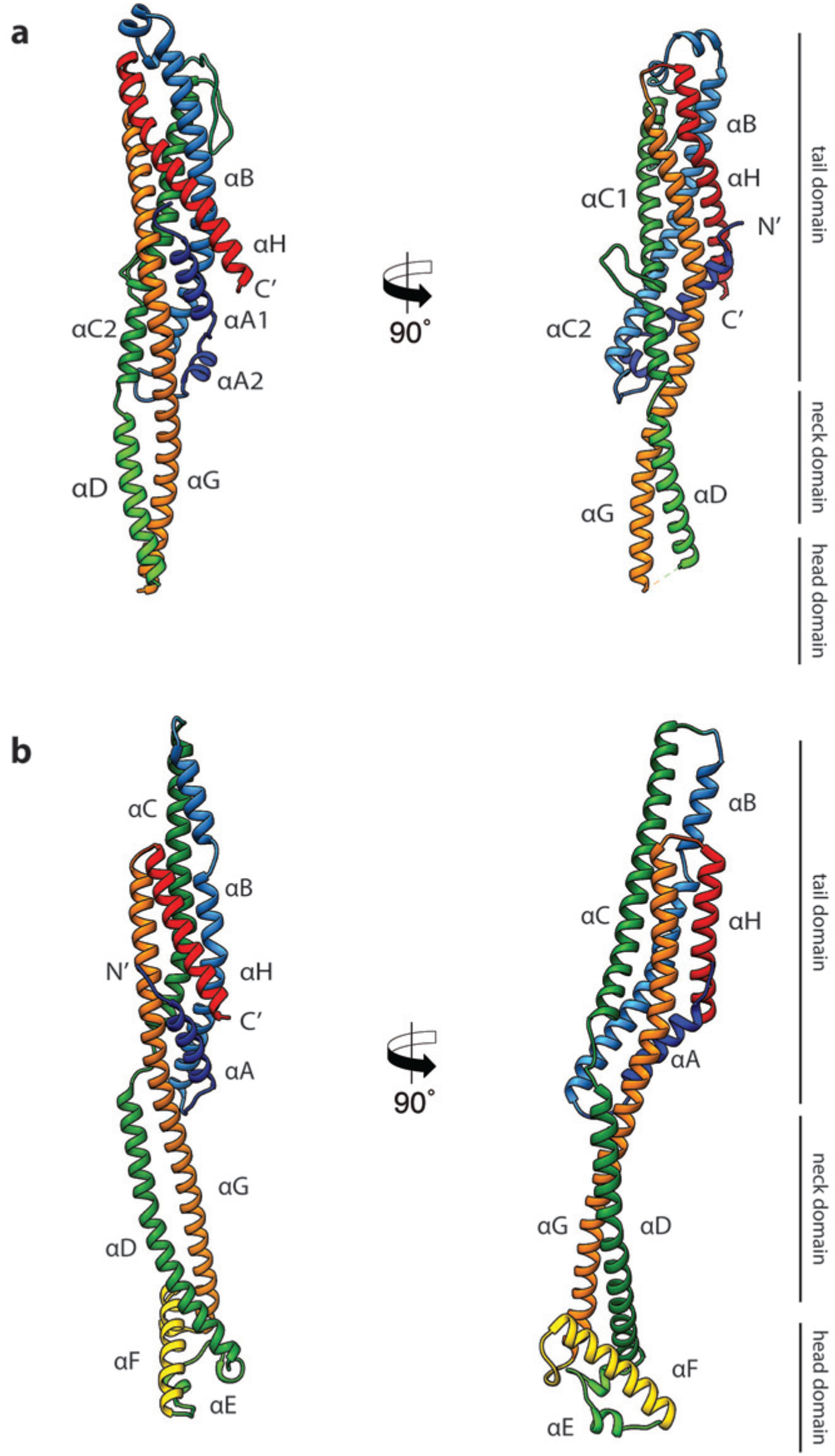
Crystal structures of XaxA and XaxB in their soluble monomeric form. **a)** Ribbon representation of the atomic model of the XaxA soluble monomer. **b)** Ribbon representation of the XaxB soluble monomer. Each helix is depicted in a different color and labeled accordingly.

A long coiled-coil structure, composed of a continuous helix (αG) and another one that is divided into three (XaxA: αC1, αC2 and αD) or two (XaxB: αC, αD) segments, forms the backbones of XaxA and XaxB. It connects all domains and forms in both XaxA and XaxB the neck and head domain. The neck domain, that is approximately 35 Å in length, does not exist in ClyA-type toxins, which are in general more compact (Figure S2a). In XaxA, the tip of the coiled-coil, predicted as hydrophobic transmembrane region is not resolved in our crystal structure, however, secondary structure predictions for this region suggest a continuation of the coiled-coil (Figure S4).

In contrast to XaxA the head domain of XaxB contains in addition to the central coiled-coil a helix-loop-helix motif, dividing helix αE and a 21-residue long amphipathic helix (αF). The highly conserved hydrophobic face of helix αF is oriented towards helices αD and αG and thereby shielded from the solvent (Figure 1b, 2). The conformation of the head domain is stabilized by conserved hydrophobic as well as electrostatic interactions, including putative salt bridges (Figure 2).

**Figure 2.**
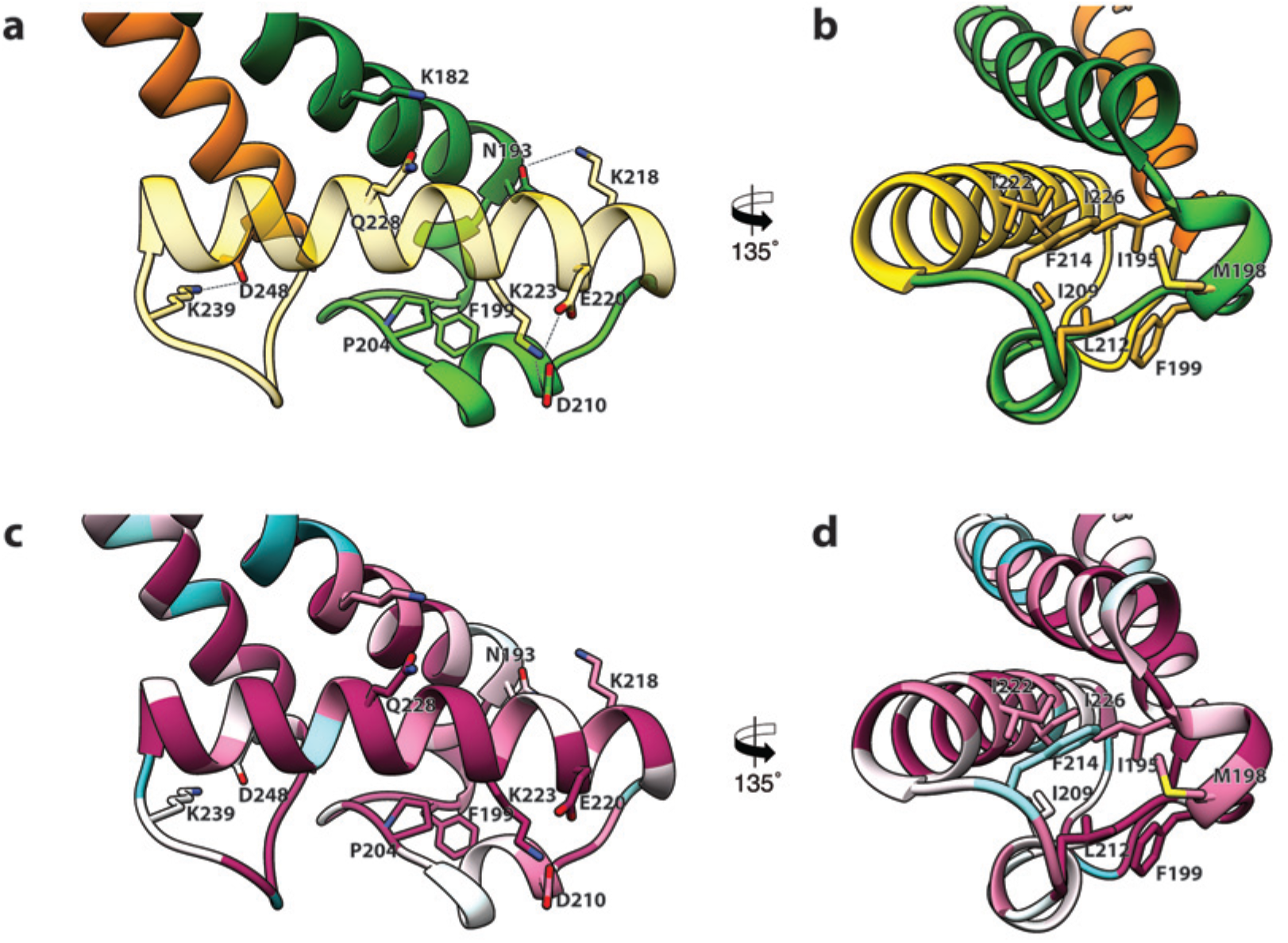
Interactions in the head domain of the XaxB monomer. **a)** The head domain of XaxB is stabilized by hydrophobic and electrostatic interactions including putative salt bridges. **b)** The hydrophobic face of the amphipathic helix αE is shielded in the soluble monomer by hydrophobic interactions with the rest of the head domain. **c,d)** Conservation of key residues in the head domain of XaxB. Same views as in (**a,b**). Figures are colored by degree of conservation based on a sequence alignment of XaxB with homologous sequences from different bacterial species from 100% (magenta) to 0% (cyan).

In general, the overall fold of the soluble monomers is similar to that of ClyA from *Escherichia coli* [20] or ClyA-type toxins, such as Cry6Aa from *Bacillus thuringiensis* [17, 18], non-hemolytic enterotoxin A (NheA) [23], and binding component B of hemolysin BL (Hbl-B) from *Bacillus cereus* [19] (Figure S2a). An important feature of ClyA and ClyA-type cytotoxins is the typical tongue motif that inserts into the membrane during pore formation [21] (Figure S2a). In ClyA, Hbl-B, NheA, and Cry6Aa the tongue is formed by a hydrophobic or amphipathic β-hairpin or a large hydrophobic loop [17, 19, 20]). Interestingly, in the case of XaxB the tongue is formed by a amphipathic helix, while XaxA does not contain such motif. Comparing the structure of XaxA with that of its pore conformation (see below) suggests that XaxA is already in its extended conformation as soluble monomer.

### Structure of the XaxAB pore complex

To investigate the pore complex formed by XaxA and XaxB, we planned to induce pore formation *in vitro* and analyze the structure of the complex by single particle electron cryomicroscopy (cryo-EM). We first mixed both soluble monomers, incubated them with a variety of detergents and analyzed the pores by negative stain electron microscopy. We could indeed observe pore formation in most cases, however, the choice of detergent greatly influenced the size and homogeneity of the observed crown-shaped pore complexes. Some detergents induced the formation of star-like aggregates or differently sized pores (Figure S5). We observed the most homogenous distribution of XaxAB pore complexes, that appear as crown-shaped structures, after incubating the monomers with 0.1 *%* Cymal-6 (Figure S5c, Figure S6c). The average diameter of the pores was ~250 Å. However, the pores had the tendency to aggregate and were not suitable for further structural investigations. Interestingly, when we incubated soluble monomers of XaxA and XaxB in the absence of detergents at room temperature, we observed the formation of higher oligomers but not of complete pores (Figure S6a,b). This indicates that dimerization and oligomerization of XaxAB can happen independently of the hydrophobic environment provided by detergents or a lipid bilayer and may happen prior to pore formation also *in vivo*.

To improve the homogeneity of our XaxAB pore complexes, we exchanged Cymal-6 with amphipols and separated the amphipol-stabilized XaxAB pores from the aggregates by size exclusion chromatography (Figure S6d,e). The thus obtained pore complexes were homogeneous and suitable for single particle cryo-EM.

Analyzing the single particles by two-dimensional clustering and sorting in SPHIRE [24, 25] revealed populations of XaxAB pores with different numbers of subunits (Figure S7). Most of the complexes contain either 12, 13, 14 or 15 subunits. We separated the different populations by multi-reference alignment and solved the structure of the different complexes in SPHIRE [25] (Methods). The average resolutions of the reconstructions were 5 Å, 4 Å, 4.2 Å and 4.3 Å for do-, tri-, tetra-, and pentadecameric pores, respectively (Figure 3, Figure S8, Figure S9). We used the highest resolved cryo-EM density of the tridecameric pore complex to build an atomic model of XaxAB (Movie S1). The high quality of the map allowed both models to be almost completely built, except for the first residues of the N-terminal helix αA in XaxA (aa 1-40) and XaxB (aa 1-12). These regions are also not resolved in the crystal structures indicating a high flexibility of the N-terminus.

**Figure 3.**
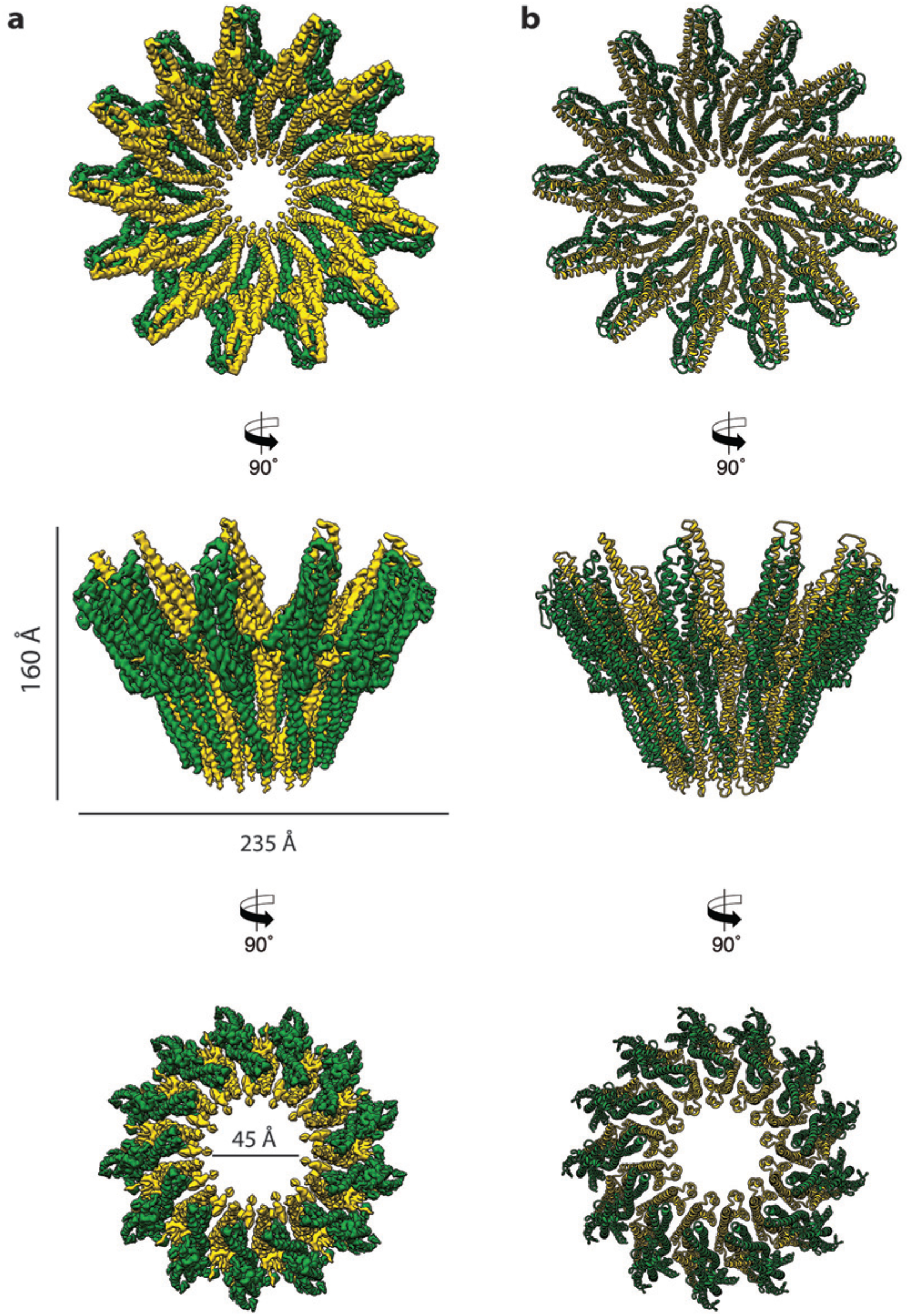
Cryo-EM structure of the tridecameric XaxAB pore complex. **a)** Cryo-EM density map of tridecameric XaxAB pores shown as top, side and bottom view. XaxA and XaxB are colored in green and yellow, respectively. **b)** Ribbon representation of the atomic model of XaxAB. Colors shown as in (**a**).

The pore complexes have a total height of 160 Å and depending on the number of subunits a diameter of 210 to 255 Å. Each subunit consists of a XaxAB heterodimer with XaxA bound to the back of XaxB. This results in a localization of XaxA on the periphery of the pore, whereas XaxB resides more at the center of the complex lining the inner pore lumen (Figure 3, Movie S1). Interestingly, the transmembrane helices of XaxA that fortify the inner ring of helices of XaxB, do not completely span the membrane (Figure S10). The arrangement of the components clearly shows that XaxAB is not a binary toxin as suggested [14, 15], but rather a bi-component toxin, such as BinAB from *Lysinibacillus sphaericus* [26, 27] and leukocidin A & B (LukGH and SF) from *Staphylococcus aureus* [28, 29] where both proteins contribute to building the pore.

Depending on the number of subunits, the inner diameter of the pore narrows down from 140-170 Å at the membrane-distal part to 40-55 Å at the transmembrane region. The inner surface of the pore is hydrophilic and mostly negatively charged suggesting a preference for positively charged ions and molecules (Figure S11). At the outside, the pore complex has a conserved highly hydrophobic band of 40 Å corresponding to the transmembrane region (Figure S11). The hydrophobic band merges into a positively charged stretch that is formed by the conserved arginine and lysine residues of XaxA (K290, K291, K293, K295, K301) (Figure S11, Figure S12). These residues likely interact with negatively charged lipid head groups of target membranes and thereby stabilize the pore complex in the lipid bilayer.

When comparing the shape of XaxAB with that of the pores of FraC and ClyA, we found that the crown-like structure of XaxA is shared by actinoporin FraC [30] but not by ClyA [21], where the extramembrane regions form a cylinder (Figure S13a). In agreement with the smaller number and size of subunits in FraC and ClyA, these pores have a smaller diameter than the XaxAB pore, and, in addition, FraC contains large β-sheets in the extramembrane region (Figure S13b). Interestingly, the lumen of all pores is negatively charged (Figure S13c), suggesting the same preference for positively charged molecules.

### Interaction between XaxA and XaxB in the pore complex

The tail domains of XaxA and XaxB do almost not differ between the oligomeric pore conformation and soluble monomers. The neck and head domains of XaxA are also arranged similarly to the crystal structure, however, the coiled-coil is twisted by 15 Å and interacts with helices αB and αC of the adjacent XaxB (Figure 4a,c, Figure 5). The neck and head domains of XaxB, however, differ considerably in comparison to the soluble monomer. The amphipathic helix αF and the helix-loop-helix motif fold out, thereby extending helices αD and αG forming the transmembrane region (Figure 4b,d, Figure 5).

**Figure 4.**
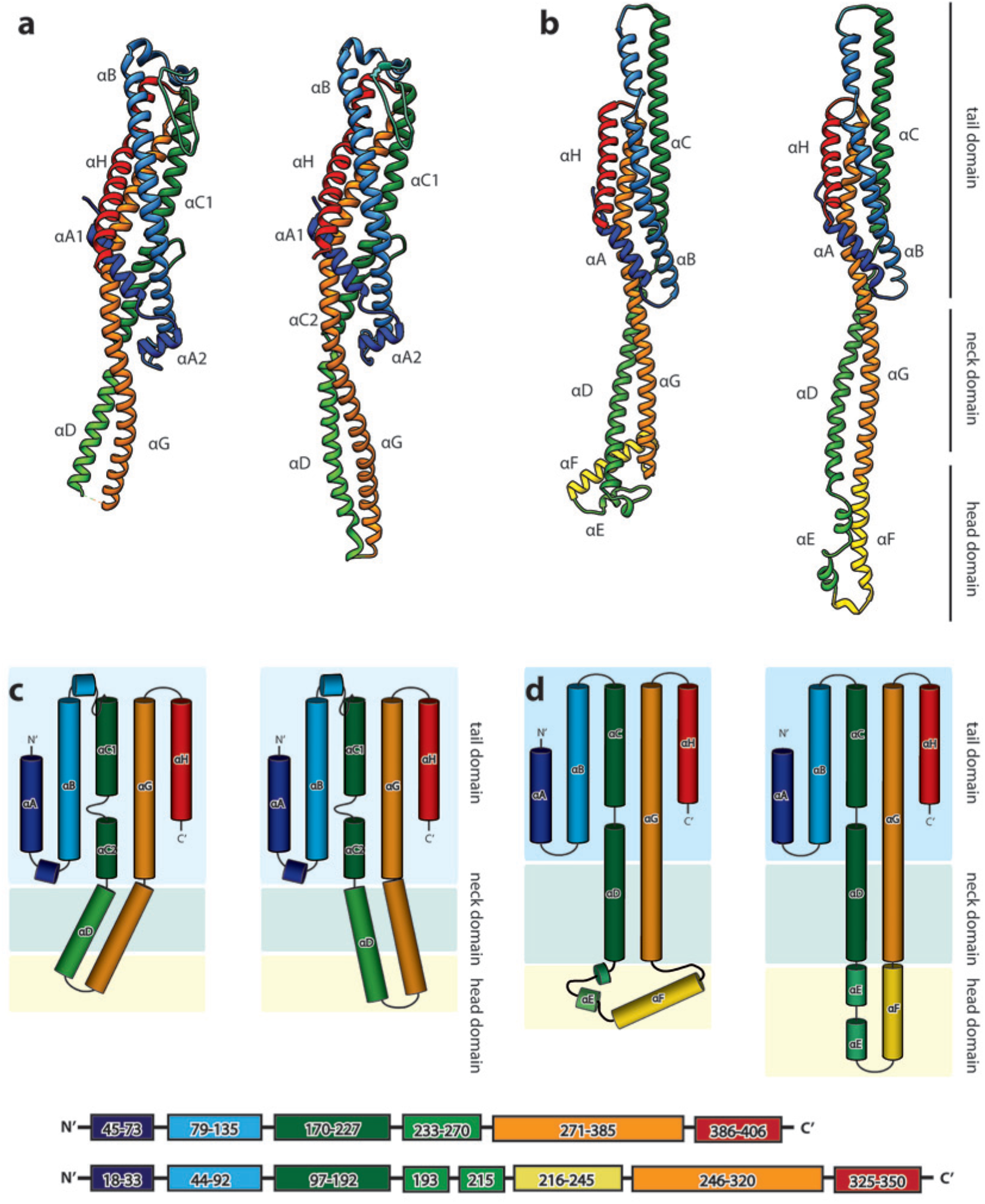
Structures of the soluble monomer and protomer of XaxA and XaxB. **a)** Ribbon representation of the atomic model of the XaxA monomer (left) and protomer (right). **b)** Ribbon representation of the XaxB monomer (left) and protomer (right). **c-d)** Topology diagram depicting helices and domain organization of XaxA (**c**) and XaxB (**d**). Each helix is shown in a different color and labeled accordingly.

**Figure 5.**
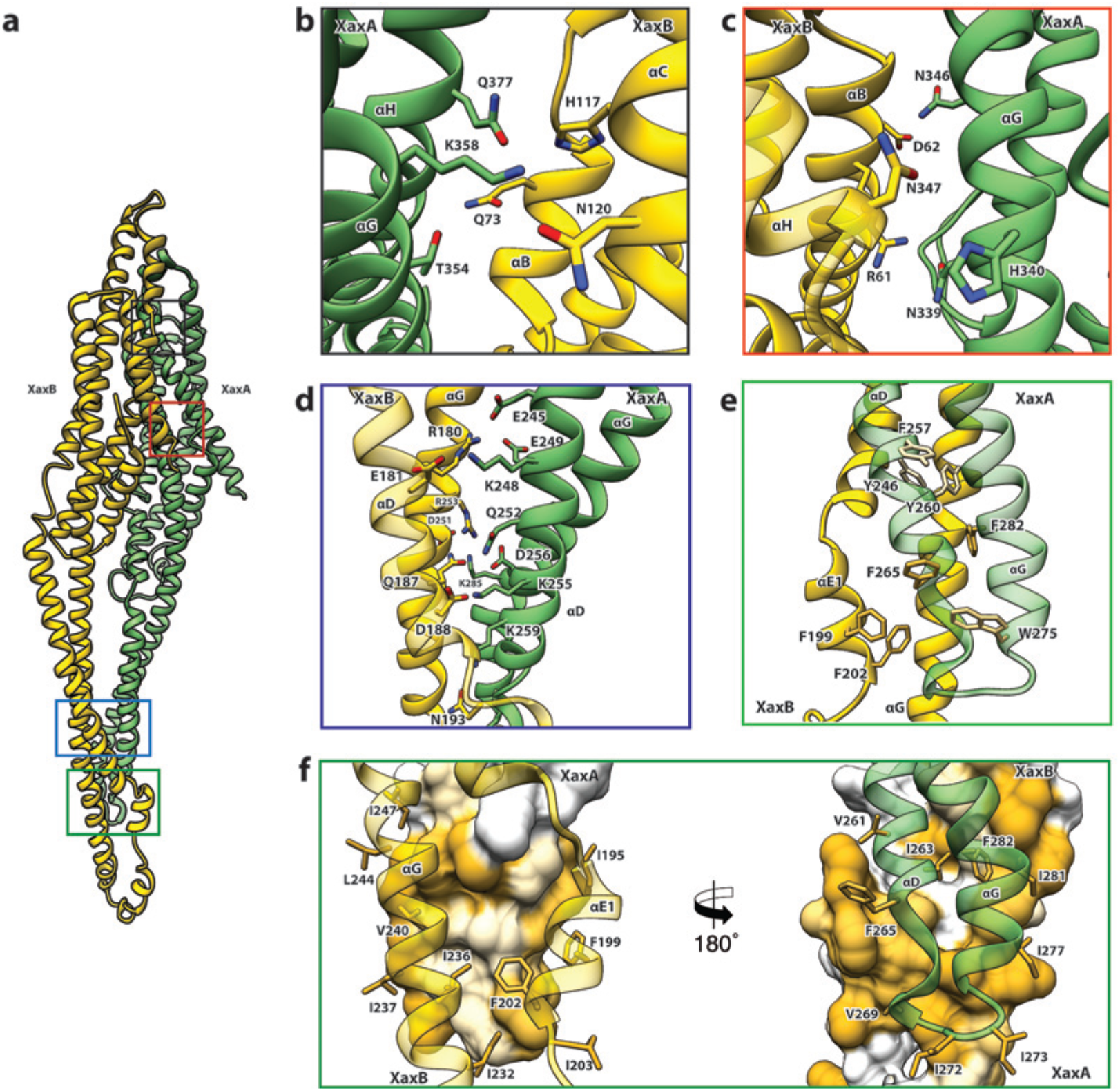
XaxAB heterodimer interactions in the pore complex. **a**) Overview of interaction interfaces between XaxA and XaxB. **b-c**) Network of putative hydrogen bonds between the tail domains. **d**) Putative salt bridges in the junction connecting the neck and head domains. **e**) The hydrophobic head domains of XaxA and XaxB are stabilized by a cluster of aromatic amino acids. **f**) Hydrophobic interface between the transmembrane domain of XaxA and XaxB in one subunit of the pore. Left: XaxB is depicted in ribbon and XaxA in surface representation colored by hydrophobicity. Right: XaxA is depicted in ribbon representation and XaxB in surface representation colored by hydrophobicity. Protomers of XaxA and XaxB are depicted in green and yellow, respectively.

The tail and head domains of XaxA and XaxB mediate interactions between the proteins in the heterodimer. We identified four major interfaces, two in the tail and two in the head domain region. The interfaces between the tail domains are stabilized by several putative hydrogen bonds and electrostatic interactions between helices αG and the C-terminal helix αH of XaxA and helices αB, αC and the C-terminal helix αH of XaxB (Figure 5a-c). Dimerization of XaxA and XaxB probably helps stabilizing the tail domain of XaxB, which took different positions in the crystal structure (Figure 5a, Figure S3).

The first interface between the head domains is formed by helices αD and αG of XaxA that interact with helices αD and αG of XaxB via a putative hydrogen network and salt bridges (Figure 5d). The second one is mediated by hydrophobic interactions between helices αF and αE of XaxB with αD and αG of XaxA (Figure 5f). A prominent feature is the high accumulation of aromatic residues at this interface (Figure 5e). Interestingly, some these residues are also involved in stabilizing the soluble XaxB monomer (Figure 2). Since most of the interfaces between XaxA and XaxB in the heterodimer locate to the tail domain and do not differ between the soluble monomer and pore conformation, we suggest that heterodimer formation precedes membrane insertion.

The heterodimers are linked manifold in the oligomeric pore. One XaxA interacts simultaneously with XaxA and XaxB of the adjacent heterodimer. The same is true for XaxB that interacts with both XaxA and XaxB of the adjacent heterodimer (Figure 6). Two major interfaces are mediated by the tail domains of XaxA and XaxB (Figure 6a-c). The residues K45, N50, E398, E402 and D333 that are conserved in XaxA form an extensive putative hydrogen network and salt bridges with helices αB (R48, Y52) and αC (D138, R147) of the adjacent XaxB (Figure 6b). A second putative hydrogen network between helices αC2 and αG in XaxA and helices αD and αG in XaxB likely contributes to the stabilization of the oligomer (Figure 6c). The oligomer is further stabilized by a putative salt bridge between two XaxAs. Glutamate E206 in the loop between αC1 and αC2 of XaxA of one subunit interacts with lysine K405 in the C-terminal helix of neighboring XaxA (Figure 6e). The fourth interface is formed by the head domains of two XaxBs via several putative hydrogen bonds (D197/S241 N192/K245) (Figure 6d). Taken together the heterodimeric subunits of the complex and the heterodimer itself are stabilized by strong interactions that guarantee a stable pore complex inside the membrane.

**Figure 6.**
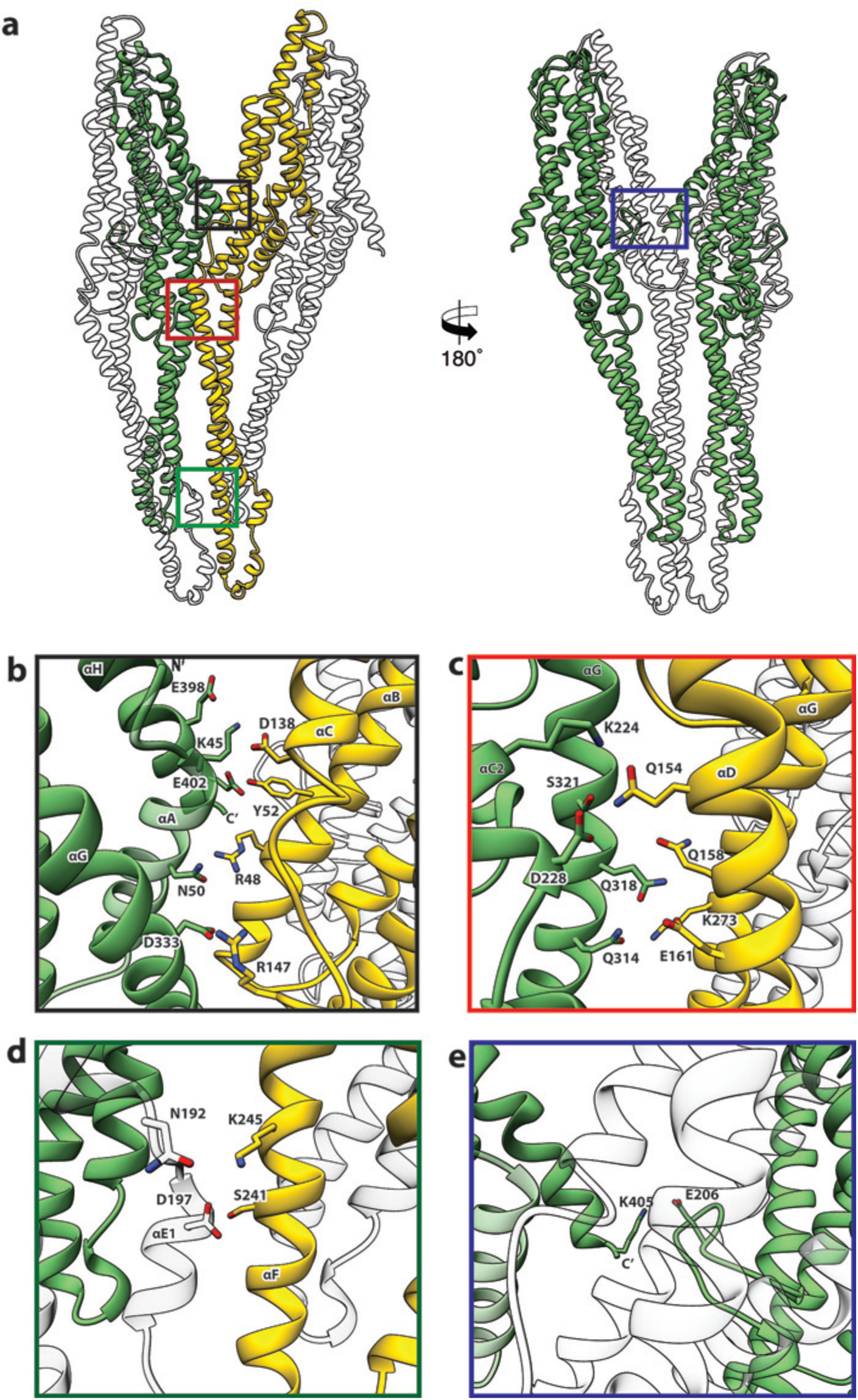
Inter-subunit interfaces of the XaxAB pore complex. **a**) Overview of four prominent inter-subunit interfaces. **b,c**) The tail domain of XaxA forms an extensive putative hydrogen network with the tail and neck domain of the adjacent XaxB. **d**) Stabilization of the transmembrane pore by additional putative hydrogen bonds and a salt bridge formed between XaxA and XaxB from the adjacent subunit. **e**) Putative salt bridge formed between the C-terminus and the loop connecting αC1 and αC2 of neighboring XaxA protomers further stabilizes the pore complex. Protomers of XaxA and XaxB are depicted in green and yellow, respectively.

### XaxAB spontaneously inserts membranes

There are at least two concerted or consecutive steps during pore formation of PFTs, namely oligomerization and membrane penetration [5]. In bi-component toxins, where both proteins contribute to building the pore, the two components first dimerize into a heterodimer prior to oligomerization [26, 28]. In several cases PFTs have been shown to oligomerize and insert spontaneously into membranes *in vitro* [2]. However, membrane insertion *in vivo* depends on the specific interaction with lipids or proteins on the membrane surface of the host [31]. To better understand the process of dimerization, membrane insertion and pore formation of XaxA and XaxB, we performed *in vitro* reconstitution assays with and without liposomes.

XaxA alone has the tendency to form small aggregates by interacting with its head domain (Figure S1). Since the short hydrophobic region of the head domain resides inside the membrane in the pore complex, we believe that the clustering of XaxA monomers is caused by mild hydrophobic interactions of these regions. This again suggests that already the soluble monomeric form of XaxA has a certain affinity to the hydrophobic environment of biomembranes. To test, whether XaxA can spontaneously insert into membranes, we incubated it with 1-palmitoyl-2-oleoyl-sn-glycero-3-phosphocholine (POPC) or brain polar lipids (BPL) liposomes and analyzed its incorporation by size exclusion chromatography and negative stain electron microscopy (Figure S14a-d). Interestingly, the protein was not incorporated into the liposomes and no larger structures could be observed on the vesicles (Figure S14c,d). This indicates that albeit the hydrophobic tip of the head domain, XaxA cannot spontaneously insert blank membranes *in vitro*. The same is true for XaxB alone. When incubated with liposomes the protein neither perforates membranes nor oligomerizes on the liposomes (Figure S14a,b,e,f).

When both XaxA and XaxB are added to liposomes, they spontaneously insert and form the typical crown-shaped pores (Figure S14g-i) as we have observed them in detergents (Figure S5e). Notably, this is independent of the sequence of mixing, i.e. XaxA can be added before XaxB or vice versa, suggesting that dimerization of XaxA and XaxB is necessary for spontaneous insertion of the proteins and subsequent pore formation. Importantly, pore formation in liposomes happens without specific lipids, such as cholesterol, or protein receptors at the membrane surface.

### Pore formation – Structural comparison between monomers and pores

In general, the transition from the soluble monomer to the protomer does not involve major structural rearrangements of the whole molecule. Only the conformation of the head domains changes considerably. Besides the described twist of XaxA (Figure 4a), the α-helical tongue αF of XaxB folds out forming the transmembrane region. Interestingly, the conformation of the coiled-coil backbone in XaxB remains unaltered (Figure 4b). This is in direct contrast to ClyA [20, 32] but similar to FraC [30], the only other two α-PFTs, for which a structure of the soluble and pore complex has been determined at high resolution. In ClyA, not only the head domain but also the tail domain undergoes considerable conformational changes [32].

In order to better understand the conformational changes during dimerization, oligomerization and pore formation, we compared the structures of the soluble and pore forms of XaxA and XaxB. When the crystal structures of XaxA and XaxB are overlaid with the respective XaxAB structure, it becomes obvious that the neck and head domains of the proteins would not interact (Movie S2). In agreement with our reconstitution assays such a dimer would probably not be able to spontaneously insert into membranes. In the XaxAB pore conformation, however, helices αD and αG of XaxA, forming the coiled-coil backbone twist by 15 Å towards XaxB (Figure 4a, Figure 5, Movie S2). As described above, through this conformational change a stronger interaction with XaxB is created. Interestingly, without the conformational change in XaxA, oligomerization of XaxAB would not be possible because of prominent steric hindrances (Movies S3). This movement is therefore crucial for complex formation.

If we assumed that only XaxA and not XaxB changed its conformation during dimerization and oligomerization (Movie S2, Movie S3), the transmembrane region of XaxA would sterically clash with the loop between αF and αG of XaxB from the adjacent subunit (Movie S2, S3). This could in principle trigger conformational changes in XaxB that activate its head domain for membrane insertion.

To better analyze these hypothetical conformational changes in detail, we created a heterodimer model comprising the cryo-EM structure of XaxA (XaxA_prot_) and the crystal structure of XaxB (XaxB_mon_) and analyzed interfaces and residues that might trigger membrane insertion (Figure 7). We identified two hinge regions that facilitate the swinging out movement of αF in XaxB (Figure 7, Movie S2, Movie S3). One hinge region is located in the hydrophobic loop between the short helices of the helix-loop-helix motif. It contains a highly conserved proline residue (P204) that is also involved in stabilization of the soluble monomer (Figure 2). The second hinge is located in the loop connecting αF and αG, including the conserved residue G243 (Figure 2, Figure 7).

**Figure 7.**
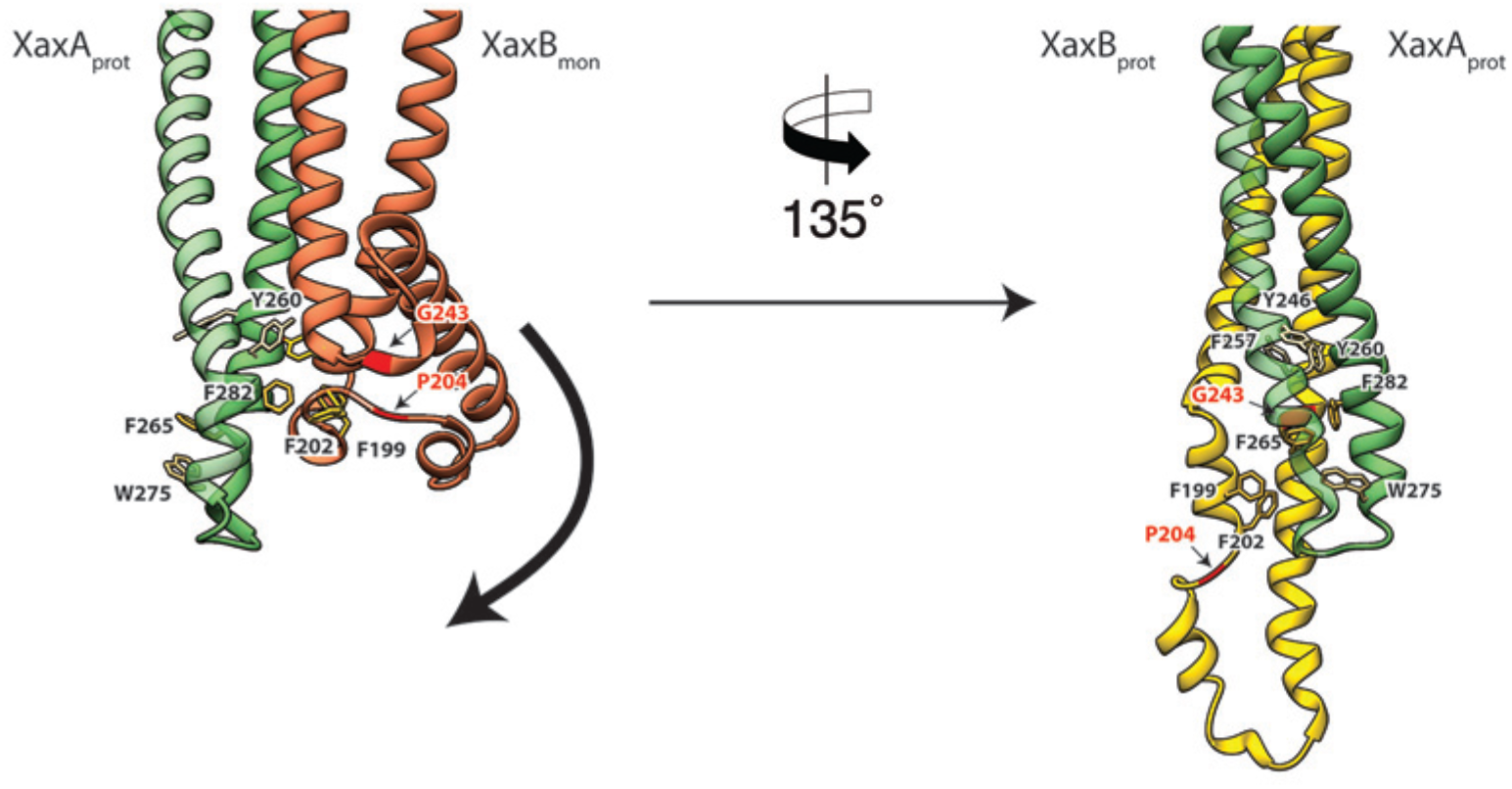
Model for membrane insertion. A heterodimer model was built with XaxA in protomeric (XaxA_prot_) and XaxB in monomeric (XaxB_mon_) conformation to mimic a possible intermediate state (left) and compared to the conformation in the pore complex (right). An aromatic cluster at the bottom of the head domain of the XaxA_prot_-XaxB_mon_ heterodimer possibly triggers the conformational change of XaxB when exposed to a lipid membrane. Swinging out of the amphipathic helix αE happens at two hinge regions at the position of conserved proline (P204) and glycine (G243) residues respectively (highlighted in red and marked with arrows). After membrane insertion, the aromatic residues interact with each other, stabilizing the new conformation.

A cluster of aromatic residues at the bottom of the head domain of our XaxA_prot_-XaxB_mon_ heterodimer model suggests that this region could be crucial in triggering the conformational changes in XaxB when exposed to a lipid membrane. In the heterodimer of the XaxAB pore complex, most of these residues build a hydrophobic cluster between the trans-membrane domain of XaxA and the reorganized helix αE of XaxB (Figure 7). Aromatic residues have been shown to be important for membrane insertion of many PFTs and responsible for conformational changes induced by their interaction with membranes or detergents [21]. Interactions with the membrane likely destabilize this region, inducing stronger conformational changes in the rest of the domain.

### Mechanism of pore formation

Our atomic model of XaxA and XaxB in solution as well as in the pore conformation provides important insights into the interaction and function of these proteins. Although the structural record is lacking intermediate states, we can use the information provided by our structural data to define critical steps in the action of XaxAB toxins and suggest the following mechanism.

Although XaxA and XaxB are not homologous, their structure is very similar. The two components of the xenorhabdolysin form heterodimers, 12 to 15 of which assemble into membrane-perforating pores. In contrast to previous predictions [14, 15], XaxAB is therefore not a typical binary toxin, but rather bi-component α-PFT. So far only structures of bi-component β-PFTs have been reported. Our structure of the XaxAB pore represents the first structure of a bi-component α-PFT.

Our results show that XaxA and XaxB together form higher oligomers in the absence of detergent or membranes. In addition, XaxA likely activates XaxB during oligomerization by inducing conformational changes. We therefore propose that XaxA and XaxB dimerize (Figure 8a-c) and oligomerize (Figure 8d) in solution. Dimerization happens probably spontaneously since the conformation of domains located at the heterodimer interface in the tail domains of XaxA and XaxB is not different compared to the monomers. The conformational change in the neck and head domain of XaxA (Figure 8b, 5d-f) further stabilizes the interaction and is crucial for oligomerization (Figure 8d). During oligomerization XaxA sterically clashes with the loop connecting helices αF and αG in XaxB. We therefore propose that XaxA induces conformational changes in XaxB that do not immediately result in exposing its hydrophobic domain but rather put XaxB in an activated state for membrane insertion (Figure 8b-c). When interacting with a lipid membrane, aromatic residues at the bottom of the head domain of XaxB likely trigger the conformational change resulting in membrane perforation (Figure 8d). Our reconstitution assays in liposomes showed that neither XaxA nor XaxB strongly interact with liposomes. Thus, neither the interaction of the aromatic residues of XaxB nor the hydrophobic domain of XaxA are able to enter membranes on their own and dimerization and induced conformational changes during oligomerization are crucial for membrane insertion. Since XaxB is the component that finally forms the pore, XaxA that only partially enters the membrane, acts like an activator of XaxB and stabilizes it in the pore complex. Obviously, more evidence is needed before our proposed mechanism of XaxAB action can be regarded as established. Thus, additional structures of intermediate states are needed to fully understand the process.

**Figure 8.**
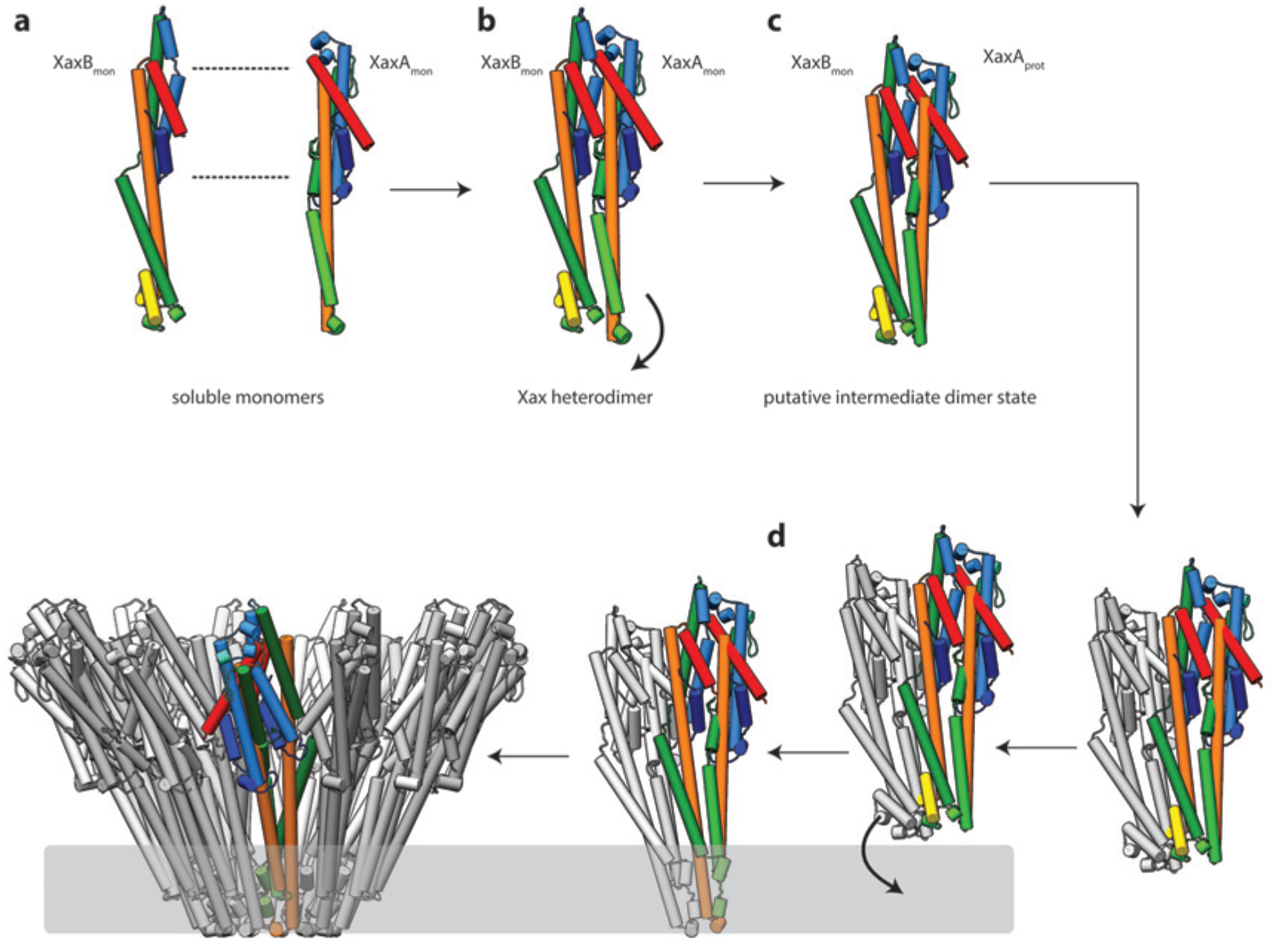
Mechanism of pore formation. **a**) XaxA and XaxB dimerize in solution. **b-c**) The major interaction site in the heterodimer is between the tail domains of XaxA and XaxB (**b**). This interaction induces of neck and head domain (αD, αG) of XaxA to shift towards XaxB (αD, αG) activating XaxB for oligomerization Interaction (**c**) and membrane insertion by clashing with the loop between αE, αF. **d**) Interactions of aromatic residues at the bottom of the head domain with the membrane trigger the conformational changes that lead to membrane insertion.

In summary, our results provide novel insights into the mechanism of action of xenorhabdolyins and serve as a strong foundation for further biochemical experiments to fully understand the molecular mechanism of xenorhadolysin intoxication.

## Methods

### Protein expression and purification

The genes coding for C-terminally His6-tagged XaxA and N-terminally His6-tagged XaxB were introduced into a pET19b vector and expressed in the *E. coli* BL21 RIPL (DE3) expression strain. Both constructs contained a PreScission cleavage site. For the expression culture, 2 l of LB media containing 125 μg/ml ampicillin were inoculated with the preculture and cells were grown at 37 °C until an OD_600_ of 0.5 – 0.8 was reached. Selenomethionine-substituted XaxB was expressed in the *E. coli* BL21 RIPL (DE3) strain in M9 minimal medium with the addition of 100 mg/l L-lysine, 100 mg/l L-phenylalanine, 100 mg/l L-threonin, 50 mg/l L-isoleucine, 50 mg/l L-leucine 50 mg/l L-valine and finally 60 mg/l l-selenomethionine (SeMet). Afterwards, protein production was induced by adding 0.4 mM of isopropyl-β-D-thiogalactopyranoside (IPTG) and incubated for 20 h at 20 °C. The cells were harvested and the bacterial pellet homogenized in a buffer containing 50 mM HEPES, pH 7.5, and 200 mM NaCl. After cell disruption, the lysate was centrifuged at 38,000 rpm, 4 °C and XaxA and XaxB was purified using Ni-NTA affinity and size-exclusion chromatography (Superdex 200 26/600, GE Healthcare).

### Crystallization of XaxA and XaxB

Crystallization experiments were performed using the sitting-drop vapor diffusion method at 20 °C. XaxA crystals formed by mixing 0.1 μl of 40 mg/ml purified XaxA with 0.1 μl reservoir solution containing 0.2 M sodium chloride, 0.1 M phosphate citrate pH 4.2 and 10 % PEG 3000 over a period of three weeks. SeMet-labeled XaxB (40 mg/ml) was mixed in a 1:1 ratio with reservoir solution containing 0.2 M NaBr, 0.1 KCl and 20 % PEG 3350 with a final drop size of 2 μl. Prior to flash freezing in liquid nitrogen, the crystals were soaked in reservoir solution containing 20 % glycerol as cryo-protectant.

### X-ray data collection and processing

X-ray diffraction data for XaxA was collected at the PXIII-X06DA beamline at the Swiss Light Source (24 ds) and at the DESY PETRA III beamline P11 (3 ds) from one crystal. The datasets were merged and used for phase determination.

Data collection for XaxB was performed at the PXII-X10SA beamline. Datasets were indexed, integrated and merged with the XDS package [33, 34].

### Structure solution and refinement

XaxA crystallized in orthorhombic space group P2_1_2_1_2_1_ with a unit cell dimension of 67 × 90 × 153 Å and two molecules per asymmetric unit (AU). Phases were determined using the anomalous sulfur signal of the merged datasets and HKL2MAP [35], the graphical interface for SHELX C/D/E [36]. The obtained phases combined with the given sequence and a few placed α-helices in the density with COOT [37] were sufficient enough for phenix autobuild [38] to almost completely build the structure of XaxA. The structure was refined with the datasets collected at the DESY PETRA III beamline P11. XaxB also crystallized in orthorhombic space group P2_1_2_1_2_1_ with a unit cell dimension of 89 × 99 × 194 Å and four molecules per AU. The diffraction data of XaxB was processed with the XDS package and SeMet atoms were determined using the CRANK2 pipeline [39, 40] in the CCP4 software package [41]. SHELX C/D [36] was used in the substructure detection process, while REFMAC [42], SOLOMON and PARROT [43]were used for phasing & substructure refinement and density modification for hand determination, respectively. BUCANEER [44] gave the best results for the initial model-building step. This model was first optimized with phenix autobuild [38]. The rest of the model was built in COOT [37] using the anomalous peaks of the SeMet residues to determine the amino acid sequence due to the limited resolution. The structures were optimized by iteration of manual and automatic refinement using COOT [37] and phenix refine implemented in the PHENIX package [45] to a final Rfree of 28 *%* and 30 *%* for XaxA and XaxB, respectively (Table S1).

### Reconstitution into liposomes

Stock solutions of 10 mg/ml 1-palmitoyl-2-oleoyl-sn-glycero-3-phosphocholine (POPC) and brain extract polar lipids (BPL) (Avanti Polar Lipids) were prepared in buffer containing 20 mM Tris-HCl pH 8, 250 mM NaCl and 5 % w/v n-octyl-β-D-glucopyranoside (Antrace). 10 μM XaxA and XaxB were mixed with a final lipid concentration of 2 mg/ml and incubated for 30 minutes at room temperature. For reconstitution, the mixture was dialyzed against a buffer containing 20 mM HEPES pH 7.5 and 200 mM NaCl. The sample was then analyzed by size exclusion chromatography with a Superose 6 10/300 GL column (GE Healthcare Life Sciences) and by negative stain electron microscopy.

### Preparation of XaxAB pore complexes

XaxAB pore complexes were prepared by incubating equimolar concentrations of XaxA and XaxB with 0.1 % cymal-6 (Antrace) at room temperature overnight. For a more homogenous and stable distribution of XaxAB pore complexes the detergent was exchanged to amphipols A8-35 (Anatrace). Amphipols were added in fivefold molar excess and the solution was incubated at room temperature for 20 minutes. For detergent removal, the mixture was dialyzed against a buffer containing 20 mM HEPES pH 7.5, 200 mM NaCl overnight at room temperature. Subsequently, aggregates and XaxAB pore complexes with higher molecular weight were separated by size exclusion chromatography on a Superose 6 10/300 GL column (GE Healthcare Life Sciences).

### EM data acquisition

The quality of the XaxAB pore complexes was evaluated by negative stain electron microscopy before proceeding to cryo-EM grid preparation. 4 μl of a 0.01 mg/ml XaxAB solution in amphipols were applied to a freshly glow-discharged copper grid (Agar Scientific; G400C) coated with a thin carbon layer and incubated for 45s. After sample incubation, the solution was blotted with Whatman no. 4 filter paper and stained with 0.75% uranyl formate. The digital micrographs were acquired with a JEOL JEM-1400 TEM equipped with an acceleration voltage of 120 kV, and a 4,000 × 4,000 CMOS detector F416 (TVIPS) with a pixel size of 1.33Å/pixel.

For sample vitrification, XaxAB pore complexes were concentrated to a final concentration of 1 mg/ml and 4 μl sample was applied onto freshly glow-discharged holey carbon grids (C-flat 2/1, Protochips), incubated for 45 s, blotted for 2.5 s and plunged into liquid ethane with a CryoPlunge3 (Cp3, Gatan) at 90 *%* humidity. The grids were then stored in liquid nitrogen.

A cryo-EM dataset of XaxAB in amphipols was collected with a Cs-corrected TITAN KRIOS electron microscope (FEI), with a XFEG and operated at an acceleration voltage of 300 kV. Images were acquired automatically using EPU (FEI) and a Falcon III (FEI) direct detector operated in counting mode at a nominal magnification of 59,000 x corresponding to a pixel size of 1.11 Å/pixel on the specimen level. In total 4,746 images were collected with 180 frames, an exposure time of 60 s resulting in a total dose of ~ 44 e^−^ Å^−2^ and a defocus range of 1.0 - 2.6 μm. Motion correction was performed using the MotionCor2 [46].

### Single particle cryo-EM data processing

All image-processing steps were carried out with the SPHIRE software package [25] (Figure S9). Initially micrographs were manually screened for bad ice or high drift and discarded accordingly. The remaining 3,617 motion-corrected sums without dose weighting were evaluated in aspect of defocus and astigmatism in CTER [25] and low quality images were discarded using the graphical CTF assessment tool in SPHIRE [25]. 186,700 single particles were automatically picked from motion-corrected sums with dose weighting using gautomatch (http://www.mrc-lmb.cam.ac.uk/kzhang/). 2-D class averages were generated as a template for gautomatch by manually picking 200 micrographs with EMAN2 boxer [47]. Pre-cleaning of the dataset and reference-free 2-D classification were performed with the iterative stable alignment and clustering approach ISAC2 [24] in SPHIRE with a pixel size of 4.97 Å/pixel on the particle level. Refined and sharpened 2-D class averages with the original pixel size and exhibiting high-resolution features were generated with the Beautifier tool implemented in SPHIRE (Figure S8b). The quality of the 2-D class averages were examined in regard of high-resolution features and completeness of the XaxAB pore complexes. According to observed oligomerization states of XaxAB pore complexes in the class averages, five initial 3-D models with c12, c13, c14, c15 and c16 symmetry were generated with RVIPER. Particles were then sorted against the five RVIPER models using the 3-D-mulrireference projection matching approach (sxmref_ali3d). The clean dataset was split into four datasets according to the number of XaxAB subunits in the complex: c12: 4,409 particles, c13: 53,546 particles, c14: 46,596 particles and c15 34,542 particles. The sixteen-fold symmetry was discarded due to low number of particles (193). The subsets containing particles with thirteen-, fourteen- and fifteen-fold symmetry were further cleaned with ISAC and subsequently subjected to 3-D refinements in MERIDIEN with a mask excluding amphipols and applying c12, c13-, c14-, and c15-symmetry, respectively [25]. In the following only the results of the map with the highest resolution will be described in detail.

SPHIRE’s PostRefiner tool was used to combine the half-maps, to apply a tight adaptive mask and a B factor of −170 Å^2^. The estimated average resolution according to the gold standard FSC@0.5/0.143 criterion between the two masked half-maps was 4.5/4 Å for the c13-symmetry (Figure S8f). The estimated accuracy of angles and shifts at the final iteration of the 3-D refinement was 0.55 degrees and 0.6 pixels, respectively. The ‘Local Resolution’ tool in SPHIRE (Figure S8e) was used to calculate and analyze the local resolution of the c13 density map. The resulting colored density map showed a local resolution of up to 3.4 Å at the lower tail domain region, whereas the tip of the spikes at the top of the XaxAB pore and at the end of the transmembrane region showed the lowest resolution (5 − 6.7 Å) (Figure S8e). The final density was locally filtered according to the estimated local resolution using the ‘LocalFilter’ tool in SPHIRE. Details related to data processing are summarized in Table S2.

### Model building, refinement and validation

The atomic model of the XaxAB pore complex was built by isolating the EM density of a XaxAB dimer and rigid body fitting the crystal structure of XaxA into the EM density map using UCSF Chimera [48]. XaxA was further fitted into the dimer density using IMODFIT [49]. For XaxA only the transmembrane region (aa 254-283) had to be manually built, which was missing in the crystal structure. The final model of the XaxA protomer covers residues 41 - 405 of the full-length sequence with residues 1 - 40 missing at the N-terminal helix αA. XaxB was built by placing helix fragments into the remaining density with COOT [37], generating first a polyalanine model and subsequently determining the correct sequence by the identification of bulky side chains. The full sequence of the XaxB protomer is also almost covered in the final model (aa 13-350) with the first 12 residues missing at the N-terminal helix αA. The XaxAB dimer was then rigid-body fitted into the XaxAB pore complex using UCSF Chimera and the full model refined using PHENIX real-space refinement [45]. Finally, the overall geometry of the refined model was evaluated with MOLPROBITY [50]. The data statistics are summarized in Table S2.

## Author contribution

S.R. designed the project. E.S. expressed, purified and crystallized XaxA and XaxB; E.S. and I.R.V. processed, refined and analyzed X-ray data, E.S. prepared cryo-EM specimens, D.P. recorded cryo-EM images. E.S. and P.A.P. processed cryo-EM data, and E.S. built the atomic models, analyzed the data and prepared figures and movies. E.S. and S.R. wrote the manuscript. All authors discussed the results and commented on the manuscript.

## Acknowledgements

The crystallographic experiments were performed on the X10SA and X06DA beamlines at the Swiss Light Source, Paul Scherrer Institut, Villigen, Switzerland and beamline P11 at PETRA III, DESY, Hamburg, Germany. We thank the X-ray community at the Max Planck Institute Dortmund and Eckhard Hofmann from the Ruhr-University Bochum for help with data collection and Vincent Olieric and Saravanan Panneerselvam for technical support and help during SAD data collection. We thank T. Wagner and C. Gatsogiannis for support in electron microscopy image processing and K. Vogel-Bachmayr for technical assistance. This work was supported by the Max Planck Society (to S.R.); the European Council under the European Union’s Seventh Framework Programme (FP7/2007–2013) (Grant 615984 to S.R.)

## Accession numbers

The electron density map after post-processing has been deposited to the EMDB under accession code XXXXX. The final model XXXX was submitted to the PDB under the accession code XXXXX.

---

**Figure S1.**
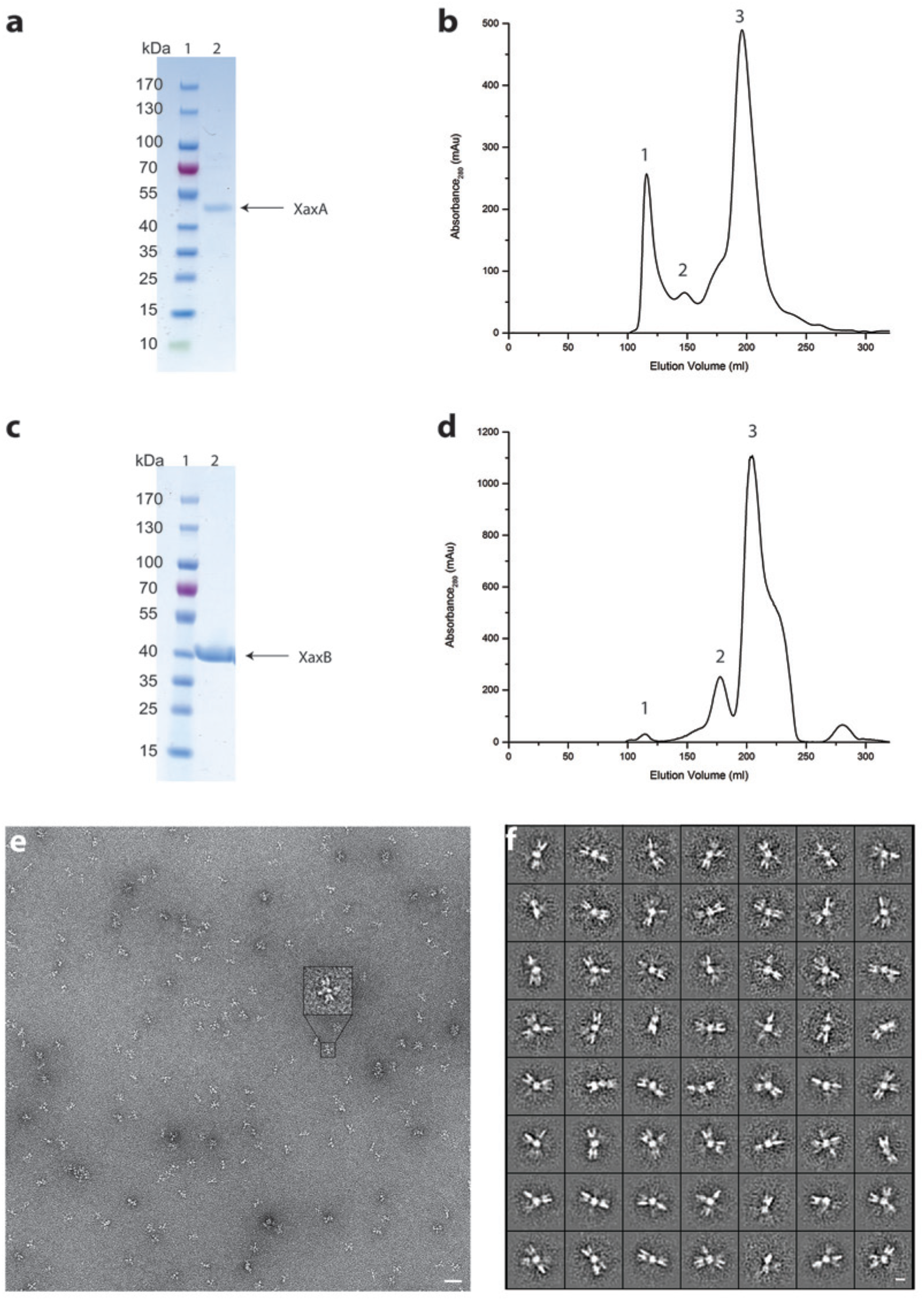
Purification of XaxA and XaxB and negative stain EM of XaxA. **a-b**) SDS-PAGE (**a**) of the peak fraction (peak 3) from the size exclusion chromatography of XaxA (**b**). **c-d**) SDS-PAGE (**c**) of the peak fraction (peak 3) from the size exclusion chromatography of XaxA (**d**). Lane 1: molecular weight marker, lane 2: protein. **e**) Representative electron micrograph of negatively stained XaxA from peak 2 in (**b**). Scale bar, 50 nm. **f**) Representative 2-D class averages of XaxA clusters. Scale bar, 10 nm.

**Figure S2.**
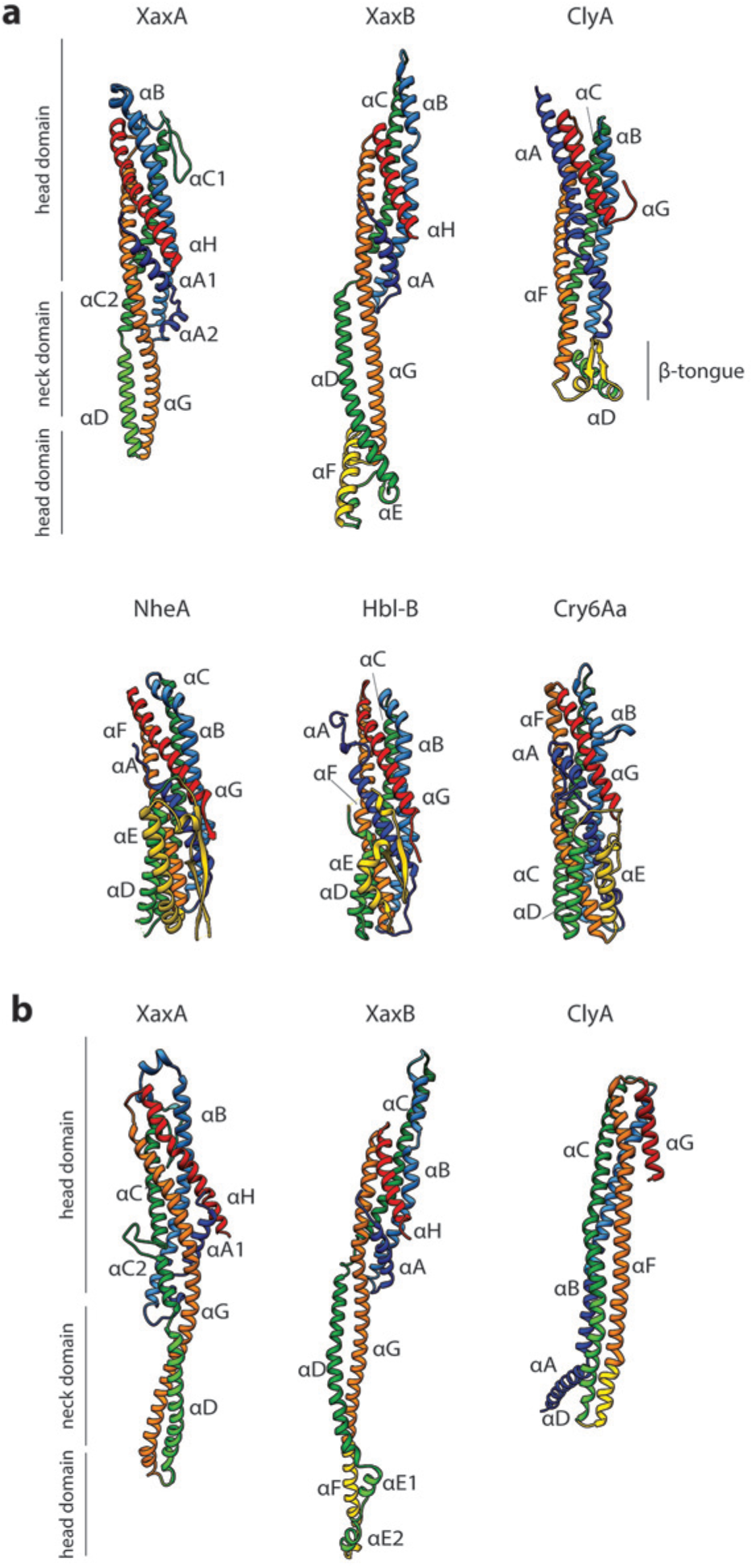
Comparison of XaxA and XaxB with ClyA-type toxins. **a**) Comparison of the soluble monomers of ClyA-type toxins (pdb-IDs: ClyA: 1QOY, NheA: 4K1P, Hbl-B: 2NRJ, Cry6Aa: 5KUC). **b**) Comparison of XaxA and XaxB protomers with the ClyA protomer (pdb-IDs: ClyA: 2WCD). Relevant helices are colored corresponding to their structural equivalent of XaxA and XaxB.

**Figure S3.**
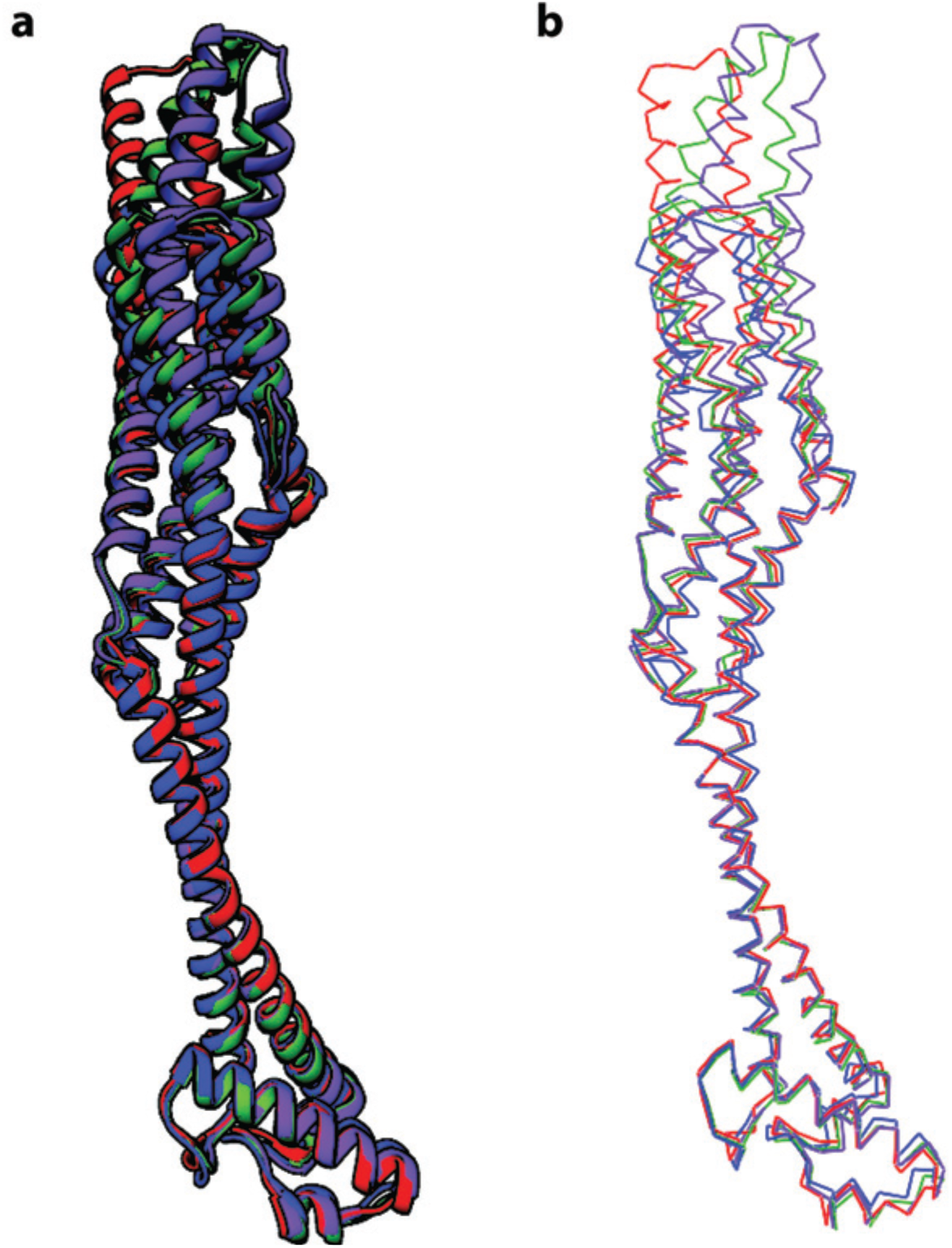
Superposition of the four XaxB molecules in the asymmetric unit. **a-b**) Molecules are depicted as ribbons (**a**) and as backbone chain trace (**b**). Molecules A, B, C, and D are shown in red, green, blue and purple, respectively.

**Figure S4.**
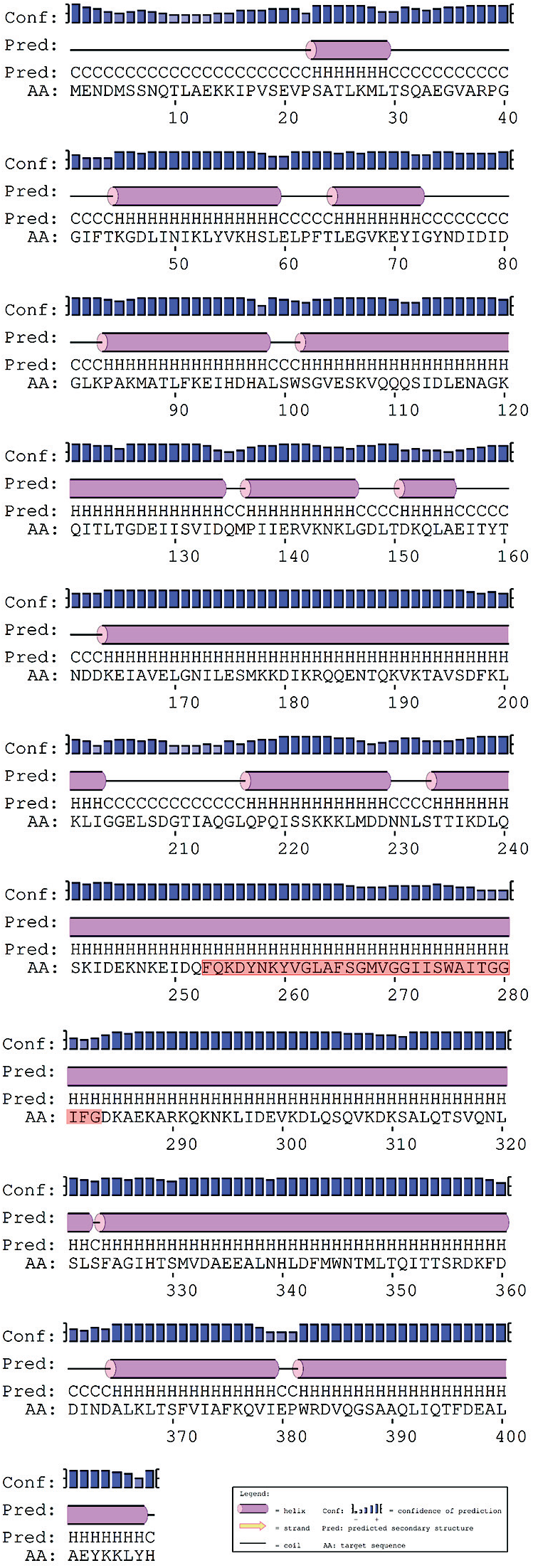
Secondary structure prediction of XaxA generated by PSIPRED. The transmembrane sequence of XaxA is highlighted in red.

**Figure S5.**
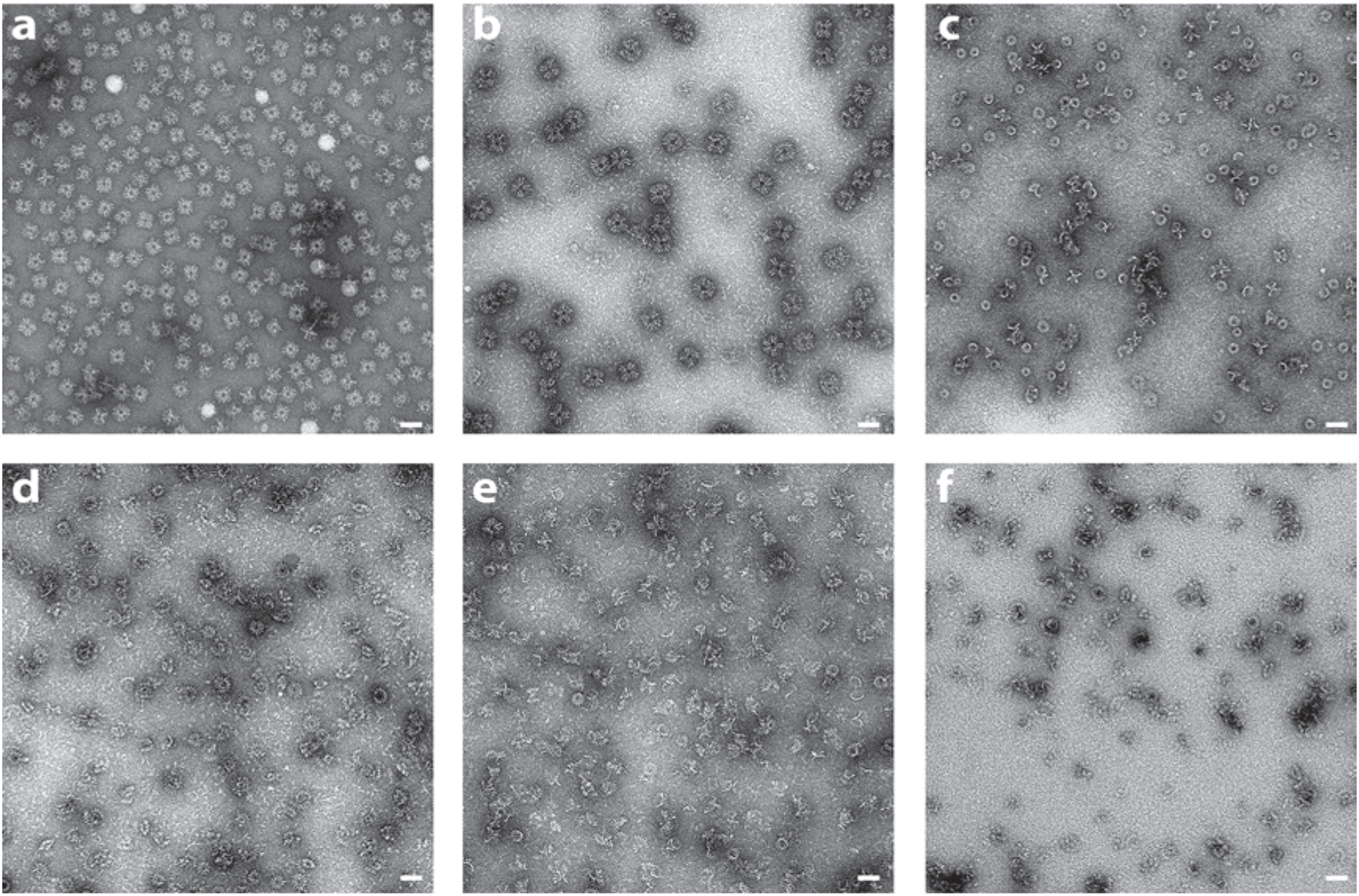
Pore formation of XaxAB induced by different detergents and analyzed by negative stain EM. **a-f**) Representative micrographs of negatively stained XaxAB pore complexes formed by incubation with different detergents, in particular n-Octyl-β-D-glucopyranoside (OG) (**a**), Lauryl maltose neopentyl glycol (LMNG) (**b**), 6-Cyclohexyl-1-hexyl-β-D-maltoside (Cymal-6) (**c**), 7-Cyclohexyl-1-hexyl-β-D-maltoside (Cymal-7) (**d**), 3-[(3-Cholamidopropyl)-dimethylammonio]-1-propane sulfonate] (CHAPS) (**e**), and digitonin (**f**). Scale bars, 50 nm.

**Figure S6.**
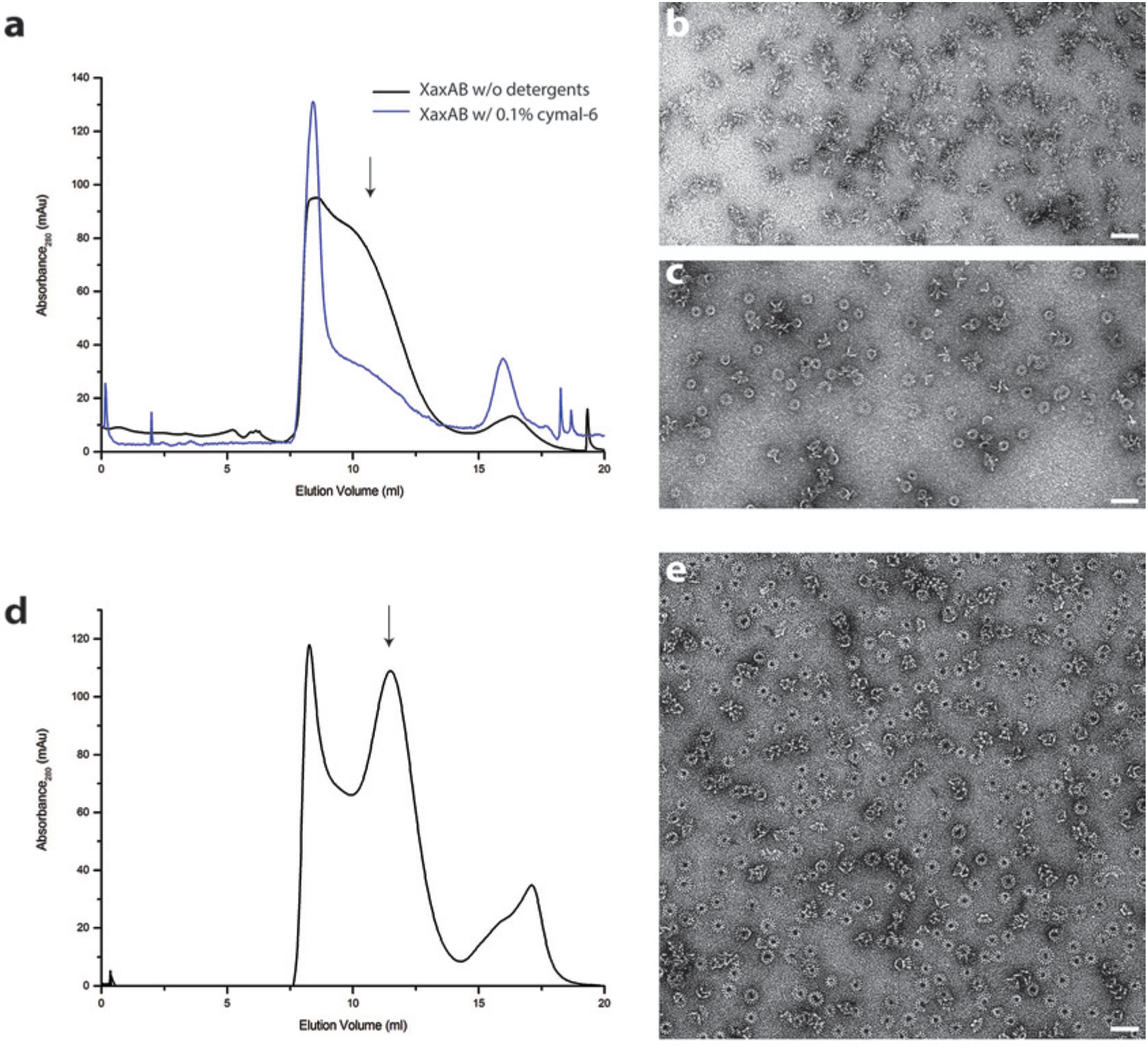
Analytical size exclusion chromatography and negative stain EM of XaxAB in Cymal-6 and amphipols. **a**) Size exclusion chromatography profiles of XaxAB after incubation overnight without (blue) and with Cymal-6 (black). **b-c**) Representative electron micrograph of negatively stained XaxAB after incubation overnight without (**b**) and with Cy-mal-6 (**c**). **d-e**) Size exclusion profile (**d**) and representative electron micrograph (**e**) of the negatively stained XaxAB pore complex in amphipols. Scale bars, 50 nm.

**Figure S7.**
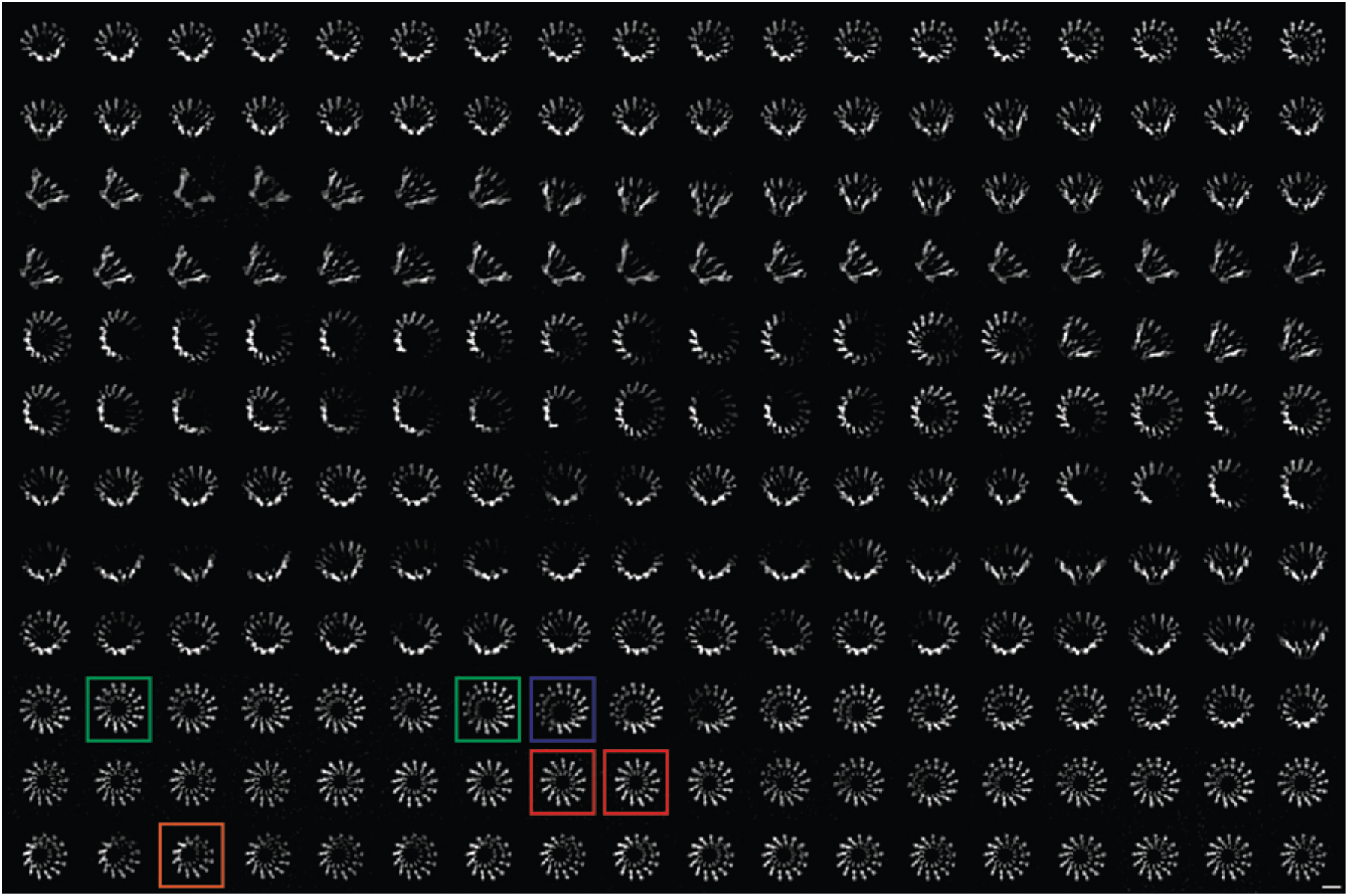
2-D class averages of XaxAB pores with different numbers of subunits. Representative class averages of top views showing XaxAB pores with 12, 13, 14, and 15 subunits are highlighted in orange, red, green and blue, respectively. Scale bar, 10 nm.

**Figure S8.**
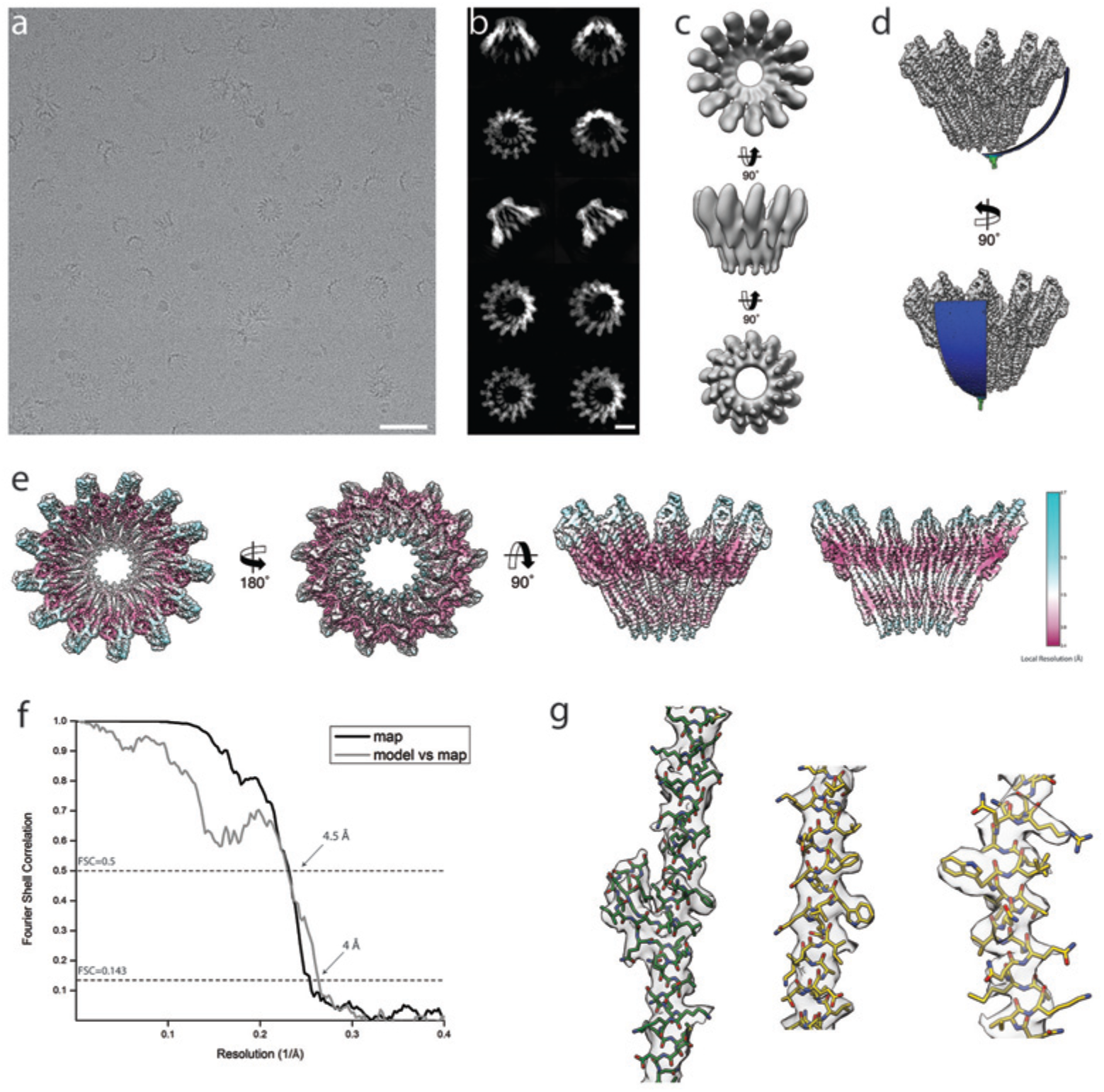
Cryo-EM structure of XaxAB. **a-b**) Representative digital micrograph (**a**) and selected 2-D class averages (**b**) of the XaxAB toxin complex embedded in vitrified ice. Scale bars, 50 nm (**a**), and 10 nm (**b**). **c**) *Ab initio* 3-D reconstruction generated with RVIPER. **d**) Angular distribution of the particles. **e**) Cryo-EM density map of XaxAB colored according to the local resolution. **f**) Fourier Shell Correlation (FSC) curve between maps from two independently refined half data sets (black) and the final map versus the atomic model (grey). The 0.143 criterion shows an average resolution of 4 Å. **g**) Representative regions of the density with fitted atomic models of XaxA (green) and XaxB (yellow)

**Figure S9.**
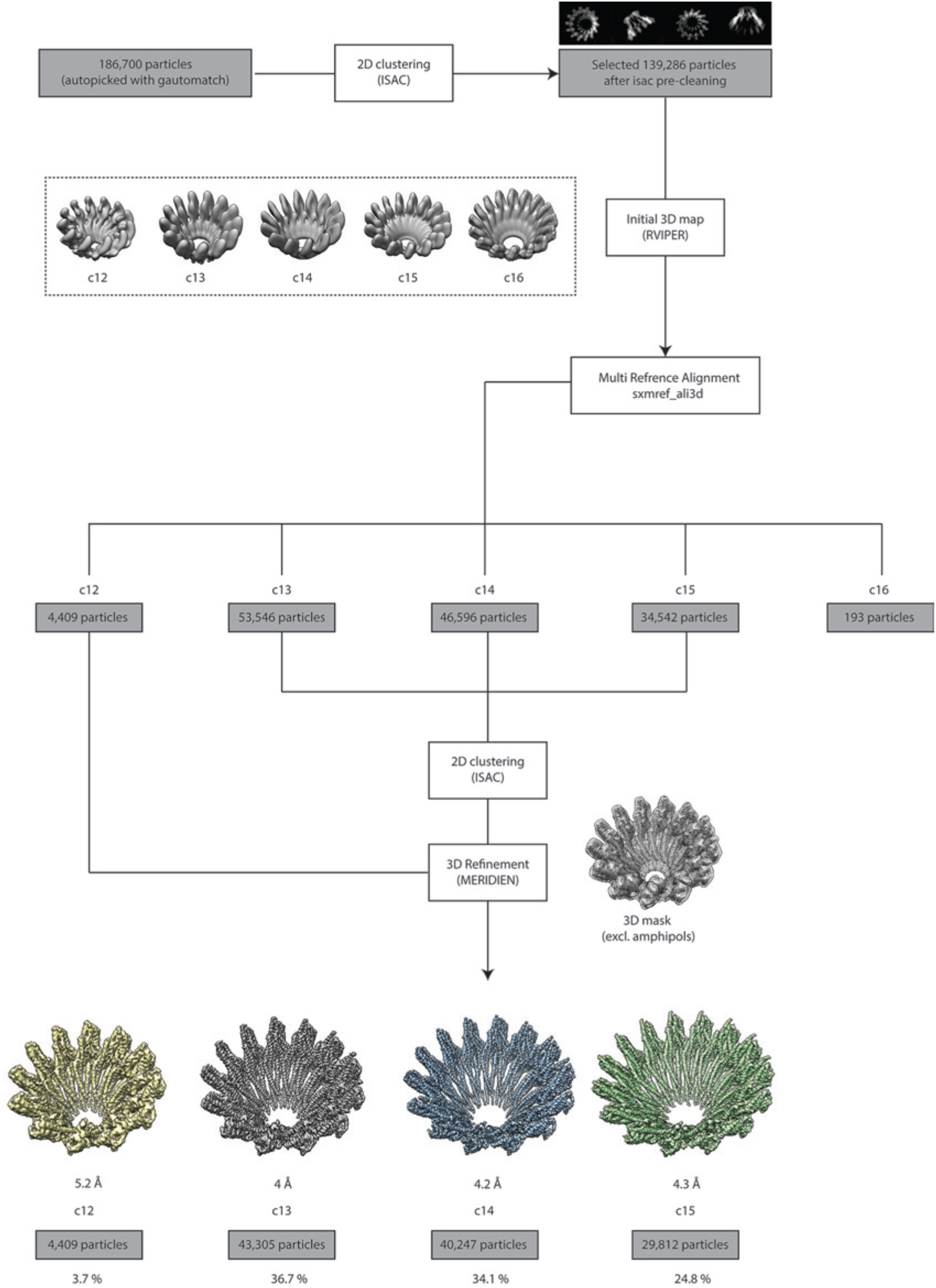
Single particle processing workflow of XaxAB structure determination.

**Figure S10.**
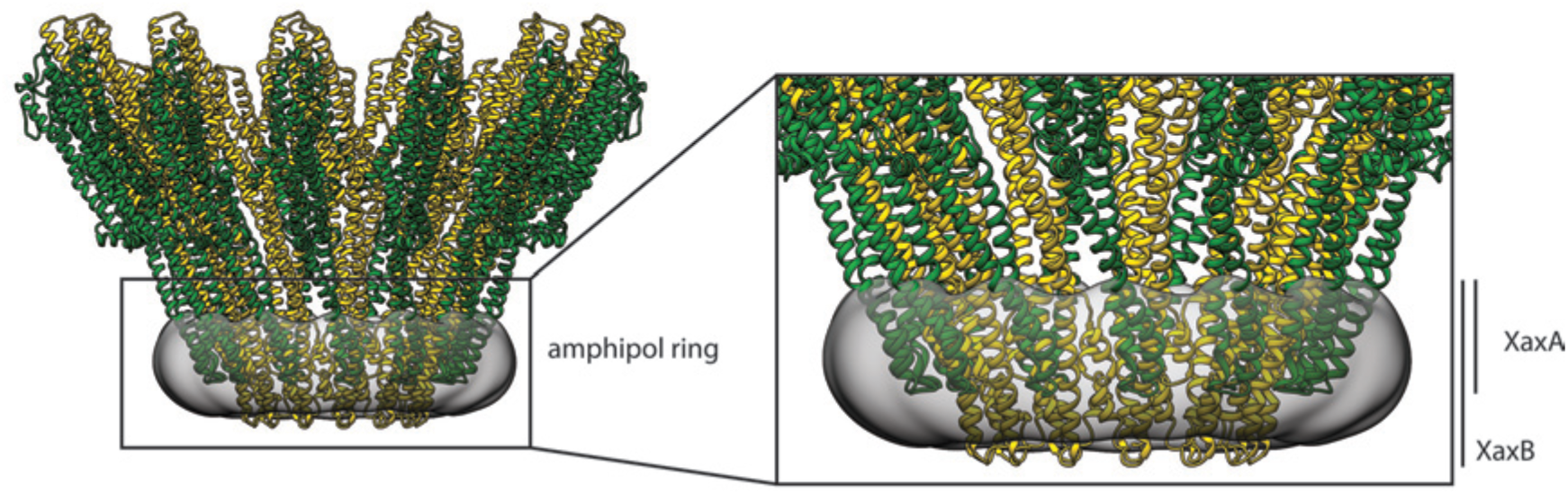
Transmembrane domains of XaxAB embedded in amphipols.

**Figure S11.**
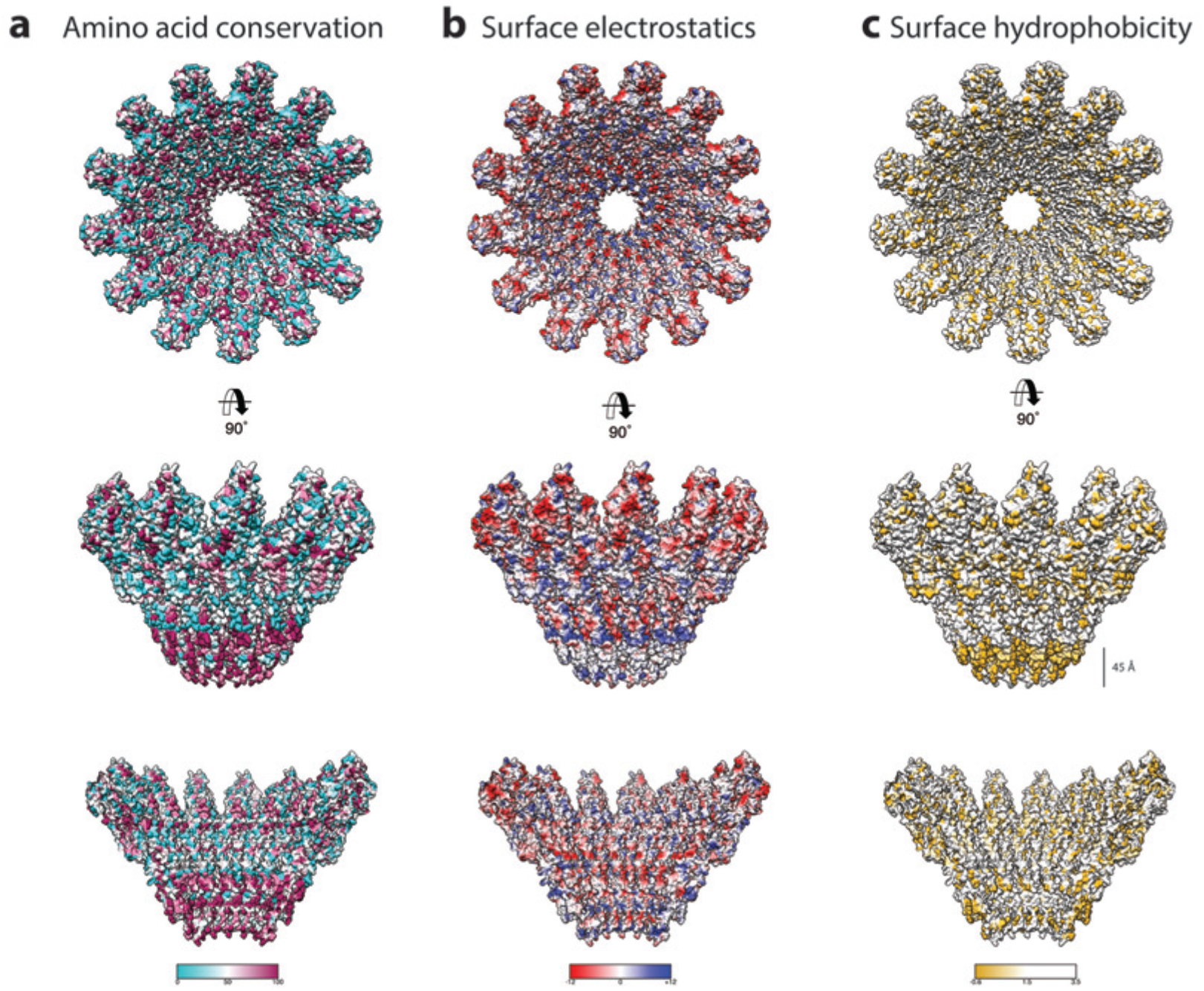
Biophysical properties of the XaxAB pore. **a-c**) Top and side views on the surface and the inside of the pore complex, showing the conservation of residues (**a**), the surface electrostatic Coulomb potential at pH 7.5 (**b**), and the surface hydrophobicity (**c**). Conserved residues are shown in magenta, positively and negatively charge surfaces are colored in blue and red, respectively and hydrophobic patches are depicted in orange.

**Figure S12.**
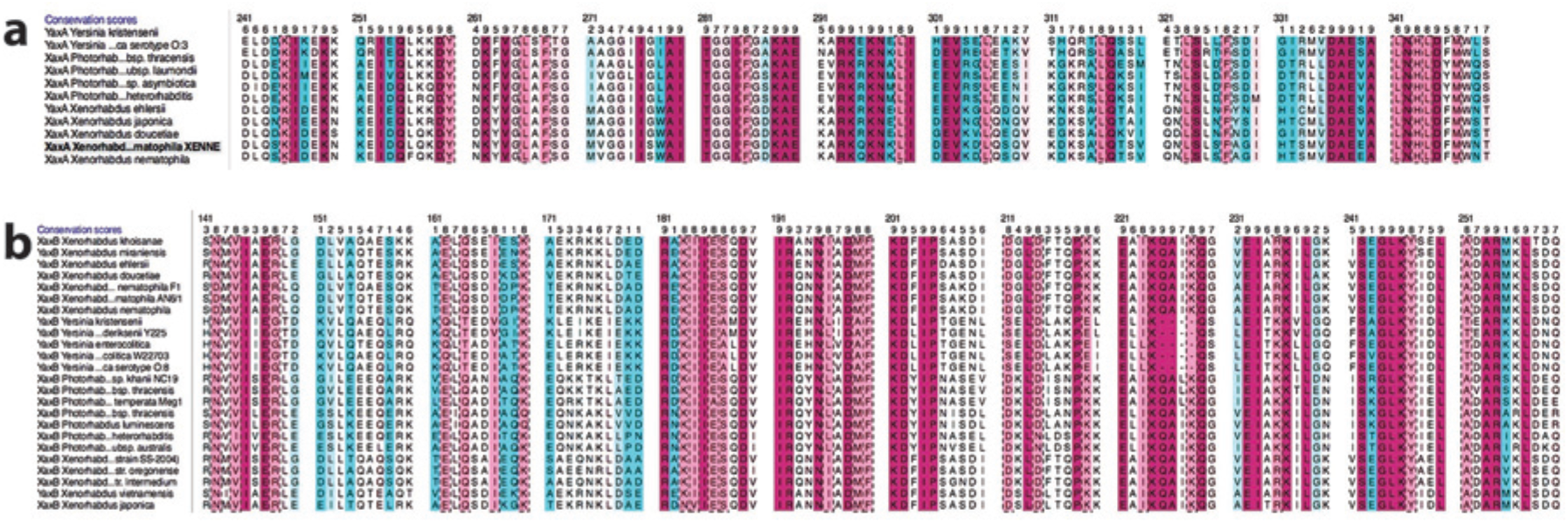
Amino acid sequence alignment and conservation of the transmembrane region of XaxA (**a**) and XaxB (**b**).

**Figure S13.**
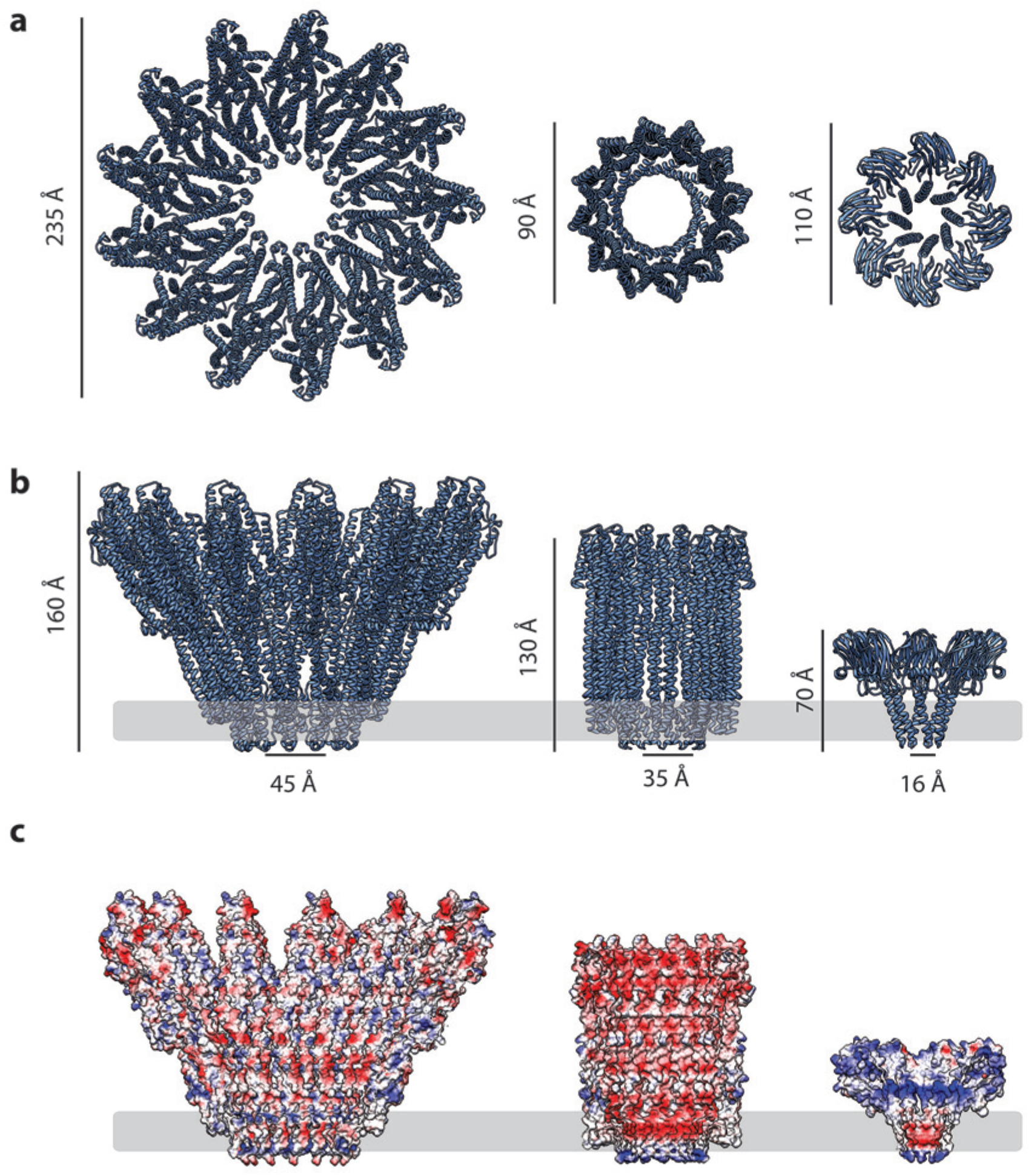
Comparison of the XaxAB pore complex with other α-PFTs. **a,b**) Top (**a**) and side (**b**) views of the XaxAB, ClyA (pdb-ID: 2WCD) and FraC (pdb-ID: 4TSY) pore complexes. **c**) Surface electrostatic Coulomb potential at pH 7.5 on the inside of the XaxAB, ClyA and FraC pore complexes. Positively and negatively charged surfaces are shown in blue and red, respectively. The membrane is indicated by a grey band.

**Figure S14.**
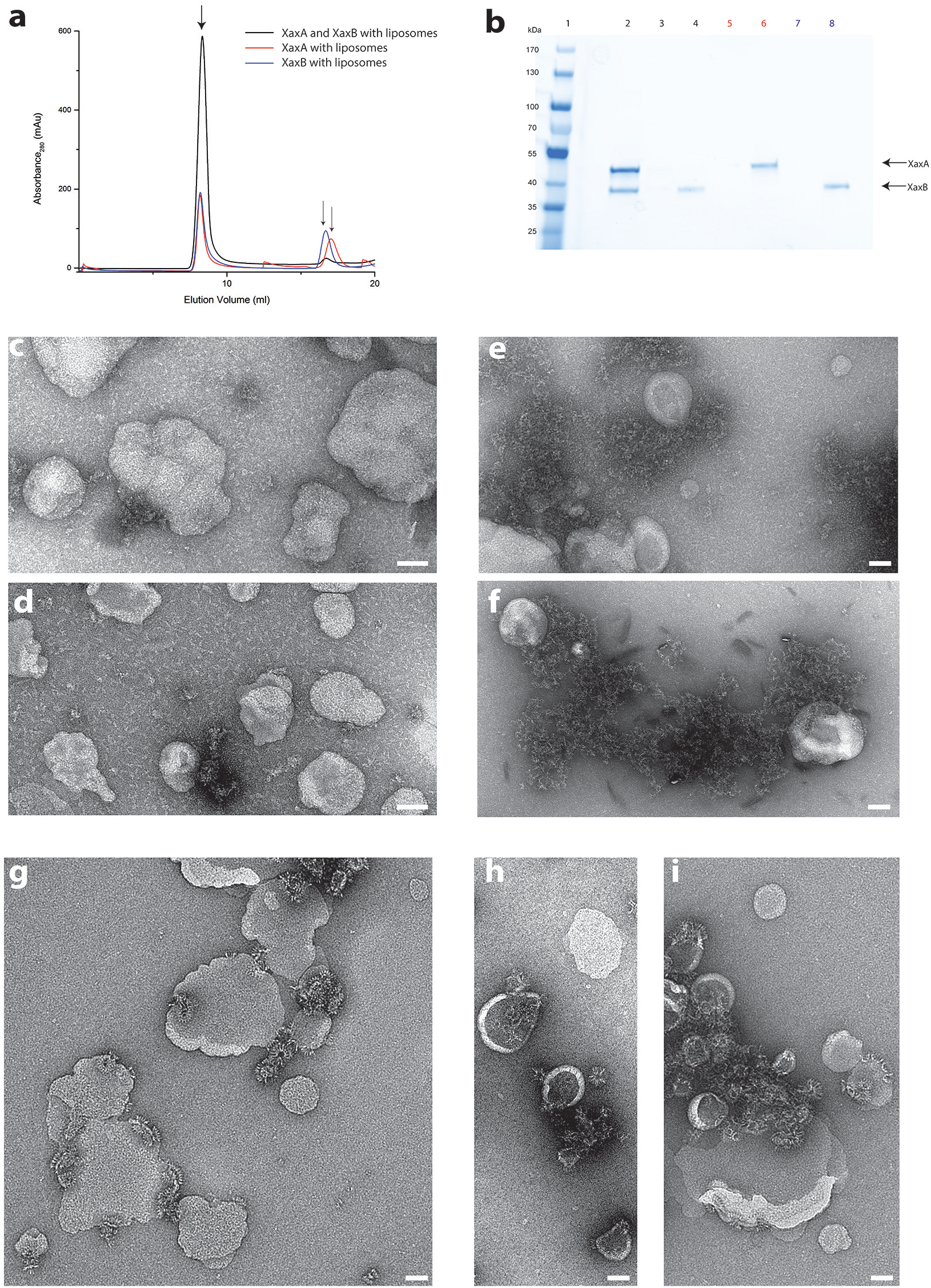
Reconstitution of XaxA and XaxB in liposomes. **a**) Size exclusion profiles of XaxA (red) and XaxB (blue) alone and of a 1:1 mixture of XaxA and XaxB (black) after incubation with liposomes. **b**) SDS-PAGE of the peak fractions of (**a**). Lane 1: molecular weight marker, lane 2-3, 4: void volume and monomeric peak of the XaxA/XaxB mixture, lane 5-6: void volume and monomer peak of XaxA, lane 7-8: void volume and monomer peak of XaxB. **c-d**) Negative stain EM of XaxA reconstitutions into POPC (**c**) or BPL (**d**) liposomes. **e-f**) Negative stain EM of XaxB reconstitutions into POPC (**e**) or BPL (**f**) liposomes. **g-i**) Negative stain EM of XaxAB reconstitutions into POPC (**g**) or BPL (**h-i**) liposomes. Scale bars 50 nm,

**Table S1.** Data collection and refinement statistics. Values for the highest resolution shell are inside brackets.

|  |  | XaxA | XaxB |
| --- | --- | --- | --- |
| Data collection |  |  |  |
| Wavelength (Å) | SLS | 2.07505 | 0.97793 |
|  | PETRA | 1.8233 |  |
| Resolution range (Å) |  | 44.48 - 2.5 (2.589 - 2.5) | 48.15 - 3.4 (3.521 - 3.4) |
| Space group |  | P 21 21 21 | P 21 21 21 |
| Cell dimensions |  |  |  |
| a, b, c (Å) |  | 67.27 90.83 153.03 | 88.7 99.41 194.15 |
| $\alpha, \beta, \gamma$ (°) | | 90 90 90 | 90 90 90 |
| Molecule no. in AU |  | 2 | 4 |
| Total reflections |  | 996,585 (92,922) | 961,813 (91,076) |
| Unique reflections |  | 33,174 (3,258) | 24,297 (2,378) |
| Multiplicity |  | 30.0 (28.5) | 39.6 (38.3) |
| Completeness (%) |  | 99.91 (99.94) | 99.91 (99.96) |
| Mean I/$\sigma$(I) | | 25.11 (2.38) | 14.23 (0.82) |
| Wilson B-factor |  | 58.45 | 137.29 |
| R-merge |  | 0.1055 (1.722) | 0.2846 (6.285) |
| R-meas |  | 0.1073 (1.753) | 0.2883 (6.369) |
| CC1/2 |  | 1 (0.872) | 0.999 (0.493) |
| CC* |  | 1 (0.965) | 1 (0.813) |
| Refinement |  |  |  |
| Reflections used in refinement |  | 33,167 (3,257) | 24,289 (2377) |
| Reflections used for R-free |  | 1,659 (173) | 1,215 (119) |
| R<sub>work</sub>/R<sub>free</sub> (%) |  | 23.84/28.57 (35.19/42.56) | 26.38/30.52 (37.37/40.11) |
| CC(work)/CC(free) |  | 0.958/0.943 (0.786/0.715) | 0.957/0.941 (0.613/0.442) |
| Average B-factor (Å<sup>2</sup>) |  | 77.47 | 142.42 |
| No. atoms in AU |  | 5,373 | 10,624 |
| macromolecules |  | 5,348 | 10,624 |
| solvent |  | 249 |  |
| Protein residues |  | 678 | 1,329 |
| r.m.s. deviations: |  |  |  |
| RMS(bonds) |  | 0.004 | 0.004 |
| RMS(angles) |  | 0.87 | 0.72 |
| Ramachandran favored (%) |  | 99.4 | 97.50 |
| Ramachandran allowed (%) |  | 0.6 | 2.20 |
| Ramachandran outliers (%) |  | 0.00 | 0.3 |
| Rotamer outliers (%) |  | 1.32 | 3.82 |
| Clashscore |  | 3.69 | 3.99 |

**Table S2.** EM Data collection and refinement statistics of XaxAB

| Data collection |  |
| --- | --- |
| Microscope | Titan Krios (Cs corrected, XFEG) |
| Voltage (kV) | 300 |
| Camera | Falcon III |
| Magnification | 59k |
| Pixel size (Å) | 1.11 |
| Number of frames | 180 |
| Total electron dose (e<sup>-</sup>/Å<sup>2</sup>) | 44 |
| Exposure time (s) | 0.88 |
| Defocus range (μm) | 1.0-2.6 |
| Number of particles | 139,286 |
| Atomic Model Composition |  |
| Non-Hydrogen atoms | 72,436 |
| Protein Residues | 9,139 |
| Refinement (Phenix) |  |
| RMSD bond | 0.006 |
| RMSD angle | 0.98 |
| Model to map fit, CC mask | 0.85 |
| Resolution (FSC@0.143, Å) | 4.0 |
| Map sharpening B-Factor (Å<sup>2</sup>) | -170 |
| Validation |  |
| Clashscore, all atoms | 4.68 |
| Poor Rotamers (%) | 0.92 |
| Favored rotamers (%) | 94.56 |
| Ramachandran outliers (%) | 0 |
| Ramachandran favored (%) | 97.42 |
| Molprobrity score | 1.35 |

**Movie SI. Cryo-EM map of XaxAB in its pore state.** Molecular model and cryo-EM map of the tridecameric XaxAB pore complex from *Xenorhabdus nematophila*, showing the overall structure of the pore complex. XaxA and XaxB subunits are colored in green and yellow, respectively.

**Movie S2. Dimerization and conformational change of XaxA and XaxB leading to the final pore complex.** Starting from the soluble monomers of XaxA and XaxB, the video focuses on the conformational changes during dimerization and membrane insertion.

**Movie S3. Interaction between the head domains of XaxA and XaxB contributes to membrane insertion of XaxB.** The video highlights possible intermediate interactions and clashes during oligomerization and membrane insertion. It starts with XaxA and XaxB in their monomeric conformation in the position of the respective protomers in the pore. Then shifts to XaxA in its pore conformation, followed by a conformational change in XaxB leading to the final XaxAB in the pore complex. Dimerization of the soluble monomers would introduce a large sterical clash between the head domains. Therefore, the soluble monomer of XaxA must transition to its protomeric form prior to oligomerization. The remaining smaller sterical clash of XaxA with the loop between helices αD, αG in the head domain of XaxB probably destabilizes its conformation and activates XaxB for membrane insertion.

